# Gluk4-containing kainate receptors regulate synaptic communication in the motor cortex and reduce axon degeneration in adult mice

**DOI:** 10.1101/2024.02.29.582867

**Authors:** Raphael Ricci, Jessica L Fletcher, Kalina Makowiecki, Renee E Pepper, Alastair Fortune, Carlie L Cullen, William M Connelly, Jac Charlesworth, Nicholas B Blackburn, Kimberley A Pitman, Kaylene M Young

**Author notes:** These authors contributed equally to this project (Jessica L Fletcher, Kalina Makowiecki). These authors contributed equally to this project (Kimberley A Pitman, Kaylene M Young).

## Abstract

Glutamate-gated kainate receptors comprising the Gluk4 subunit (encoded by *Grik4*) are highly expressed by neurons in the central nervous system. We report that *Grik4* mRNA is widely expressed by neurons in the adult mouse motor cortex, where GluK4-containing kainate receptors account for ∼60% of the kainate evoked current in layer V pyramidal neurons. To elucidate their role in motor circuit regulation, we analysed the behaviour of mice that lacked the pore forming domain of the GluK4 subunit (*Grik4^-/-^*mice). *Grik4^-/-^* mice were hyperactive, had an abnormal gait, and impaired motor coordination. At postnatal day (P)60, layer V pyramidal neurons received fewer miniature excitatory post synaptic currents, had a reduced density of thin spines on their basal dendrites, and a reduced density of VGlut1 puncta at the soma, consistent with neurons receiving fewer excitatory synaptic connections. *Grik4^-/-^* mice also lost ∼44% of their callosal axons between P60 and P180 and the amplitude of the callosal compound action potential was reduced by ∼25-30%. RNA sequencing data support the capacity for *Grik4* to modulate synaptic and neuroprotective signalling pathways.

## Introduction

The physiological role of kainate receptor signalling is poorly understood, but can change with age and differ between brain region and neuron type, being affected by the cellular localisation and subunit composition of the receptors, the presence of other interacting proteins, and whether signalling is ionotropic and/or metabotropic (for reviews see Falcon-Moya and Rodriguez-Moreno, 2021; Lauri *et al*., 2021; Lerma and Marques, 2013; Mulle and Crepel, 2021; Nair *et al*., 2021; Negrete-Diaz *et al*., 2022). Kainate receptors are tetrameric, assembled from subunits GluK1-5 (previously referred to as GluR5-7, KA1, and KA2, respectively) which are encoded by the *Grik1-5* genes. GluK4 is a high affinity subunit and requires one of the principal subunits (GluK1-3) to assemble to form a functional receptor. *Grik4* is highly expressed in the rodent hippocampus (Friedman *et al*., 1994; Wisden and Seeburg, 1993) where Gluk4-containing kainate receptors account for ∼80% of kainate receptors on CA3 pyramidal neurons (Fernandes *et al*., 2009). Hippocampal GluK4-containing kainate receptors localise to pre- and post-synaptic structures (Contractor *et al*., 2003; Mulle *et al*., 1998), mediate the kainate receptor component of excitatory post synaptic currents (Fernandes *et al*., 2009) and modulate long-term potentiation in CA3 neurons (Catches *et al*., 2012).

GluK4 is reportedly expressed at much lower levels outside of the hippocampus, however, the behavioural and pathological effects of altered Gluk4 signalling suggest that it regulates a range of CNS circuits (Aller *et al*., 2015; Arora *et al*., 2018). *In situ* hybridisation and RNA sequencing studies in mice suggest that *Grik4* mRNA is also expressed in the cerebellum (Falcon-Moya *et al*., 2018), amygdala (Arora et al., 2018) and cortex (Arora *et al*., 2018; Zhang *et al*., 2014). Genetic variants identified in *GRIK4* in humans have been associated with schizophrenia (Hu *et al*., 2022), depression and its treatment (Minelli *et al*., 2016; Pu *et al*., 2020; Ren *et al*., 2017), bipolar disorder (Koromina *et al*., 2019), and autism (Griswold *et al*., 2012). Methylation changes in *GRIK4* also implicate GluK4-kainite receptor signalling with the progression of motor dysfunction in Huntington’s disease (Lu *et al*., 2020). In mice, the germline disruption of GluK4-kainate receptor signalling results in hyperactivity, reduced anxiety, reduced depressive-like behaviour, and impaired sensorimotor gating (Catches *et al*., 2012; Lowry *et al*., 2013). By contrast, the germline overexpression of Gluk4 in the mouse forebrain (CaMKII promoter) enhances anxiety- and depressive-like behaviour in adult mice and impairs social interaction (Aller *et al*., 2015). Additional research is required to elucidate the requirement of GluK4-containing kainate receptors for non-hippocampal circuit function, particularly cortical circuit function.

Herein, we show that *Grik4* mRNA, is highly expressed throughout the motor cortex and in motor regions of the spinal cord of adult mice. While young adult *Grik4* functional knockout (*Grik4*^-/-^) mice were hyperactive, they also had abnormal gait and deficits in motor coordination that worsened with age. Layer V pyramidal neurons in the primary motor cortex had a reduced number of basal dendritic spines, particularly thin spines, and received fewer synaptic inputs. It is possible that this precipitates axon degeneration, as callosal axon density was significantly reduced by P180, resulting in a reduction in callosal compound action potential amplitude in *Grik4*^-/-^ mice. We identified several differentially expressed genes in the hippocampus of *Grik4^-/-^*mice that could provide insight into the observed changes. These data implicate Gluk4-containing kainate receptors in motor circuit maintenance in adult mice.

## Materials and methods

### Mice

All animal experiments were approved by the University of Tasmania Animal Ethics Committee (A0016151 and A0018606) and carried out in accordance with the Australian code of practice for the care and use of animals for scientific purposes. Reporting of these experiments follows the Animal Research: Reporting of In Vivo Experiments (ARRIVE) guidelines. All transgenic mice were maintained on a C57BL/6J background. Homozygous *Grik4* germline functional knock out mice (*Grik4^-/-^*; (Fernandes *et al*., 2009) were a kind gift from Professor Anis Contractor (Northwestern University). *Grik4^-/-^* were genotyped using genomic DNA that was extracted from ear biopsies by ethanol precipitation. The *Grik4* transgene was detected by PCR (Catches *et al*., 2012; Fernandes *et al*., 2009) using 50-100ng of genomic DNA and three primers: *Grik4* P1 3’ GCAGGCTGAACTCTGAGTTT, *Grik4* P2 5’ CCAGAGACAGCACTAGGTGC and *Grik4* P6 5’ GCCTGGGCTAGAGTGAGAC. *Thy1-YFPH* transgenic mice [B6.Cg-Tg(Thy1-YFP-H)16Jrs/J] were purchased from Jackson Laboratories (#003782). The YFP transgene was detected by quantitative PCR using two primers: CACCCTCGTGACCACCTT (forward) and GGTAGCGGGCGAAGCA (reverse) (Transnetyx, Memphis). Mice are referred to as wild type (WT) when they are not *Grik4^-/-^*mice and do not carry any other transgene. Mice are referred to as control (Ctrl) when they are not *Grik4^-/-^* mice but carry another transgene e.g. *Thy1-YFPH*.

Male and female mice were used equally for experiments (174 mice in total) and were assigned to experimental groups based on genotype. Where relevant, care was taken to ensure littermates were represented across experimental groups. Mice were housed in same sex groups (2-4 per cage), in individually ventilated cages (Optimice®) on a 12 h light-dark cycle (twilight phase starts 06:30, full lights on 07:00) at 21 ± 2°C with *ad libitum* access to standard rodent chow (Barrastock rat and mouse pellets) and water. However, a small cohort of C57BL/6 mice were transferred to a diet of 0.2% (w/w) cuprizone powder (C9012, Sigma) in crushed mouse food pellets (Barrestock), reconstituted with water (as per Auderset *et al*., 2020) for 5 weeks to induce demyelination and robustly induce the expression of proteins associated with axonal injury.

### Western blot protein quantification

To generate whole brain protein lysates from P5 C57BL/6 mice or *Grik4^-/-^*mice, pups were decapitated, and the brain rapidly dissected on ice to be snap frozen on liquid nitrogen and stored at - 80°C. Frozen brains were transferred into ice cold lysis buffer [50 mM Tris-HCL, 150 mM NaCl, 1% (v/v) NP-40, 1% (v/v) sodium deoxycholate, 0.1% (v/v) SDS with protease inhibitor cocktail (Roche, 1183615300)] and homogenized with a plastic pestle in a 1.5 mL tube. To generate hippocampal protein lysates, P60 mice were killed by cervical dislocation, the brains were rapidly removed and sectioned into 2 mm slices that were snap frozen on liquid nitrogen, and stored at -80°C. Hippocampi were dissected from slices spanning ∼Bregma -1.5 to -1.7 mm, placed in ice cold lysis buffer containing the protease inhibitor cocktail, and lysed using a Dounce homogenizer. Lysates were incubated on ice for 15 min then centrifuged with a benchtop centrifuge at 13,000 rpm for 15 min at 4°C and the supernatant stored at -80°C. Protein concentration was quantified by performing a modified Lowry assay (DC Protein Assay, BioRad, 5000111) with bovine serum albumin (BSA; Sigma A7030) standards according to the manufacturer’s instructions and analysed using a FLUOstar OPTIMA microplate reader at 750 nm. Protein concentration was plotted against absorbance for each standard and used to calculate the protein concentration of each sample.

The expression of each protein of interest was quantified by Western blot as previously described (Auderset et al., 2016). Briefly, 50 µg of total protein in sample buffer (4x Bolt LDS, Invitrogen, B007) containing 500 mM DL-Dithiothoreitol (DTT; Sigma, D0632) was denatured under reducing conditions by incubating at 70J for 10 min. Protein ladder (Invitrogen, LC5925 or BioRad, 1610373) or lysate was loaded into each well of a 4-12% Bis-Tris gel (Invitrogen, NW04120BOX) in MES (Invitrogen, B0002) or MOPS (Invitrogen, B000102) running buffer and the gel run at 200 V for 20 min. The protein was transferred from the gel onto polyvinylidene fluoride (PVDF) membrane (Bio-Rad, 162-0177) in transfer buffer (Invitrogen, BT0006) with 10% methanol and 0.1% Bolt antioxidant (Invitrogen, BT0005) by performing electrophoresis at 20 V for 60 min. PVDF membranes were blocked with 5% (w/v) powdered milk in 0.05% (v/v) Tween-20 in tris-buffered saline (TBS-T) then incubated overnight at 4°C with one of the following primary antibodies: rabbit anti-GluK4 (Abcam; ab10101, 1:500), rabbit anti αB-crystallin (Enzo Life Sciences; ADI-SPA-223, 1:1000), rabbit anti-KCHN3 (Invitrogen; PA5-106705, 1:1000) or rabbit anti-K_ir_2.3 (Alomone Labs, APC-032, 1:200). PVDF membranes were washed thrice in TBS-T before being incubated with goat anti-mouse-HRP (Dako, P0447) or goat anti-rabbit-HRP (Dako, P0448)] diluted 1:10,000 in 1% (w/v) powdered milk in TBS-T, for 1 hour at 37°C. Antibody labelling was detected using a chemiluminescence system (ECL; Millipore, WBKLS0500) and imaged using an Amersham imager 600 (GE Healthcare, Chicago, IL). Membranes were stripped using Restore™ PLUS Western Blot Stripping Buffer (Thermo Scientific, I464300) according to the manufacturer’s instructions and re-probed with mouse anti-β-actin (Sigma, A1978, 1:1000) or rabbit anti-GAPDH (Millipore, ABS16, 1:1000). The optical density value for each band was determined using FIJI/Image J and normalized to β-actin or GAPDH.

### Behavioural Assessments

Behavioural assessments were carried out during the dark-phase of the 12-hour light/dark cycle, as previously described (Ferreira *et al*., 2021). On the day of testing, mice were moved to the assessment room 2-hours prior to the light cycle change and habituated in the dark for 2-3 hours. All trials were video recorded and animal movement tracked using automated tracking software (EthoVision XT 11, Noldus, Netherlands). Equipment was cleaned with 70% ethanol between trials.

*Y-maze:* Black and white spatial cues were placed on all four walls of the testing arena. Lighting was set to 70 lux yellow. The polyvinyl chloride (PVC) Y-maze was configured to have an open start arm, one closed and one open arm (40cm x 10cm x 20cm), and soiled bedding from the mouse’s home cage was mixed with fresh bedding (Puracob coarse enrichment bedding, Able Scientific) and evenly placed throughout each arm of the maze. The individual mouse was placed in the start arm of the Y-maze and left for 15 min; returned to the home cage for 1 hour, and then returned to the maze – which was reconfigured so that all arms were open. The mouse was allowed to explore freely for 5 min and the distance travelled, velocity (cm/s) and time spent in the novel arm, start arm and other arm of the maze was calculated using EthoVision XT software automated tracking.

*Elevated plus maze:* The plus-shaped, PVC maze with 2 opposing open (50 cm long) and 2 opposing closed-in arms (50 cm long; 30 cm high), positioned 100 cm off the ground, and illuminated with focused light at 120 lux. An individual mouse was placed in the centre of the maze facing the open arms and allowed to explore freely for 5 min before being returned to the home cage. EthoVision XT software automated tracking was used to analyse the video to calculate the distance travelled, the velocity (cm/s) of movement, and the time spent in the open and closed arms of the maze.

*Forced swim test:* Individual mice were placed in a 20 cm diameter beaker containing ∼20 cm of ∼21°C water. Mice were closely watched for 5 min and video recorded for manual scoring by a blinded experimenter. Mice were removed, dried, and returned to their home cage, which was placed on a heat mat for 15 min after testing. Videos were scored manually to determine the proportion of time spent actively swimming or being immobile.

*Marble bury test:* Individual mice were placed in clear Optimice^TM^ cages containing 10 cm of clean bedding and 10 marbles. The marbles were evenly spaced on the surface of the bedding. Mice were left in the cage with the marbles for 30 min before being returned to their home cage. The number of marbles buried (defined as >50% of the marble height covered) was quantified.

*Open field test:* At P60, individual mice were placed in an open square PVC arena (30 cm^2^) with 20 cm walls. The light (200 lux) was focused on the centre of the arena to create a darker perimeter ∼5 cm from each edge. The mice were free to explore the arena for 10 min before being returned to their home cage. Etho Vision XT software automated tracking was used to calculate the distance travelled and velocity (cm/s) from video recordings.

*Grip strength:* P45-P180 mice underwent grip strength testing using a Grip Strength Meter (GPM-100; Melquest, Toyama, Japan). After zeroing the force gauge, each mouse was allowed to grasp the bar mounted on the force gauge. The mouse’s tail was slowly pulled back until the mouse released its paws from the bar and the tension was recorded. The test was performed in triplicate for each mouse.

*DigiGait^TM^ analysis:* Gait analysis was performed using the DigiGait^TM^ treadmill imaging system (Mouse Specifics, Inc) at P45 and reassessed at P60, P90, P120 and P180. Mice were habituated to the treadmill enclosure for 5 min before the treadmill was turned on and the speed of the transparent belt increased from 18 cm/s to 22 cm/s over a 2 min period. Short ∼10s videos were recorded from below at a belt speed of 22cm/s. A ∼3-4s excerpt, covering a time when the mouse was running straight without obvious acceleration or deceleration, was analysed using the semi-automated DigiGait^TM^ analysis software. Each digital analysis output was evaluated to correct processing errors (eg. Software mislabelled a forepaw as a hind paw) before data were exported for statistical analysis. Data for the left and right limbs were pooled, but forelimb and hindlimb data were analysed separately.

*Beam walk task:* Balance and coordination were assessed using the beam walk task at P45 and again at P60, P90, P120 and P180. A separate cohort of mice used for RNA sequencing was assessed after P300. Mice were placed at the end of a 10 mm diameter 100 cm long wood dowel. After a training session, to acclimate mice to the apparatus, mice were video recorded as they traversed the beam, entering their home cage bedding on the far side platform. Videos were used to manually score foot slips i.e. mice placed any paw below the level of the beam. A separate RNA sequencing mouse cohort was analysed at a single timepoint (>P300), using the same protocol.

### Immunohistochemistry and confocal microscopy

To obtain tissue for histological analyses, mice were transcardially perfused with 4% (w/v) paraformaldehyde (PFA; Sigma) in phosphate buffered saline (PBS); brains were dissected, and 2 mm-thick coronal slices generated using a 1 mm brain matrix (Kent Scientific). Brain slices were immersion fixed in 4% PFA at ∼21°C for 90 min; cryoprotected in 20% (w/v) sucrose (Sigma) in PBS at 4°C overnight, and snap frozen in OCT (ThermoFisher) using liquid nitrogen, before being stored at -80°C.

30 µm coronal brain cryosections, containing the primary motor cortex (M1) and underlying corpus callosum (∼Bregma +0.5), were collected and processed as floating sections (Cullen *et al*., 2019). Sections were incubated overnight at 4°C in PBS blocking solution [0.1% (v/v) Triton X-100 and 10% foetal calf serum in PBS] containing primary antibodies including: mouse anti-NeuN (1:200; Millipore, #MAB337); mouse anti-parvalbumin (1:1000, Swant, #235), rabbit anti-somatostatin (1:1000, Immunostar, #20067), mouse anti-SMI32 (1:1000, Covance/Biolegend, #801702), mouse anti-SMI312 (1:1000, Covance/Biolegend, #837904), rabbit anti-amyloid precursor protein (APP, C-terminal, 1:1000, Sigma, #A8717), rat anti-green fluorescent protein (GFP, 1:2000, Nacalai Tesque, #04404-84), mouse anti-vesicular glutamate transporter 1 (VGlut1, 1:200, Synaptic Systems, #135 511), and chicken anti-vesicular GABA transporter (VGAT, 1:1000, Synaptic Systems, #131 006). Sections were washed thrice in PBS before being incubated overnight at 4°C in PBS blocking solution containing donkey anti-rabbit (1:1000), donkey anti-mouse (1:1000), donkey anti-chicken (1:1000) or donkey anti-rat (1:500) conjugated to AlexaFluor -488, -568 or -647 (Invitrogen). Nuclei were labelled using Hoechst 33342 (1:1000; Invitrogen). Sections were washed thrice, transferred onto glass microscope slides, allowed to dry and cover slipped with fluorescent mounting medium (Dako).

Confocal images of fluorescent immunohistochemistry were collected using a spinning disk UltraView Nikon Ti Microscope with Volocity Software (Perkin Elmer) and standard excitation and emission filters for DAPI, FITC (AlexaFluor-488), TRITC (AlexaFluor-568) and CY5 (AlexaFluor-647). To identify and quantify individual cells within a defined region confocal z-stacks with 2 μm z-spacing were acquired with a 20x objective; images were collected spanning the primary motor cortex (M1) or corpus callosum (CC) and the maximum projection images were stitched together by the Volocity Software to produce a single composite image spanning the region of interest. For NeuN staining images collected at a single z-plane were stitched to create an image spanning the region of interest. To image and quantify the density of VGlut1^+^ or VGAT^+^ presynaptic sites on the soma of Layer V pyramidal neurons in the M1, YFP^+^ soma were identified in Layer V, positioned within the 60x field of view, and a confocal z-stack collected with 0.5 μm z-spacing (0.23 µm/pixel).

Image analysis was carried out by researchers blind to experimental treatment. Cell number was determined by opening the images in Photoshop (v21, Adobe) or Image J (v1.52s, NIH) and performing manual counts. Cells were only quantified if they contained a Hoechst^+^ nuclei. To quantify the expression of SMI-312, SMI-32, APP or neuronal marker (NeuN), we performed image segmentation in Image J (Collins *et al*., 2019; O’Mara *et al*., 2017) and determined the proportion of the pixels within the area that were immunopositive. To quantify the density of VGlut1 and VGAT presynaptic sites associated with individual YFP^+^ soma, we used Imaris ×64 image analysis software (v9.2.0, Bitplane, Zürich, Switzerland). Z-stack images, each containing an entire YFP^+^ soma, underwent 3D rendering using the “surface” function to determine the soma volume. VGlut1 or VGAT positive puncta were analysed using the “spots” function with XY size parameters set to 0.8 (VGlut1) or 0.5 (VGAT) µm, and the z parameter set to 1 µm. Spots of VGlut1 or VGAT labelling that were close to the rendered cell soma were quantified using the “find spots close to surface” extension (Matlab R2019a, MathWorks, Sydney, Australia), with a threshold of 1. The number of spots (puncta) close to the cell surface was expressed relative to the volume of the rendered soma (number of spots/cell volume).

### Dendritic spine density and morphology

Coronal brain cryosections were collected as detailed above (see immunohistochemistry and microscopy), transferred onto slides, allowed to dry and mounted using Prolong Glass Antifade mounting medium (Thermo Fisher Scientific) and high precision coverslips (No.1.5H, Marienfeld Superior). Confocal images of YFP^+^ neurons (*Thy1-YFPH*) in layer V and their associated dendrites in the motor cortex were captured using an Olympus FLUOVIEW FV3000 (BX63LF) laser scanning microscope (514 nm excitation) with a 100x objective (Olympus Super Apochromat UPLSAPO-100XO/1.4) and z-step size 0.09-1.2 µm. Images were 1024x1024 pixels at 0.07 µm/pixel and 0.2 µs/pixel dwell time, averaged 3x by frame. Images were converted to tiff stacks for dendritic reconstruction and analysis in Neurolucida 360 (v.2021.1.3, MBF Bioscience). Dendrite segments were selected for imaging and reconstruction if the: (i) associated cell soma was identifiable; (ii) the basal dendrite was relatively flat (XY plane) and > 10µm in length, and (iii) the dendrite was sufficiently isolated from surrounding YFP^+^ dendrites, making it possible to identify individual spines. Dendrite segments were traced using the user-guided tool followed by manual adjustment. Parameters for manual spine detection were set at a minimum of 10 voxels, a minimum height of 0.3 µm, and a maximum distance of 2.5 µm from the dendrite surface, and detector sensitivity adjusted for each spine to accurately fill spine volume. Classification into thin, mushroom, and stubby morphological subtypes was performed within Neurolucida 360 using the multi-dimensional classifier with settings: head-to-neck ratio of 1.1, length-to-head ratio of 2.3, and mushroom head size of at least 0.3 µm. Manual classification was required for ∼10% of spines. For these spines, head diameter was measured across the widest portion, and the head-neck appearance evaluated. Tracings were batch analysed in Neurolucida Explorer (v.2020.3.1) and aggregated by individual dendrite to determine spine density and the proportion of spines classified as each subtype. ≥5 dendrites were analysed per mouse and the data presented as the average per mouse.

### Transmission electron microscopy

P60 and P180 mice were transcardially perfused with Karnovsky’s fixative (2.5% glutaraldehyde, 2% PFA, 0.25mM CaCl_2_, 0.5mM MgCl_2_ in 0.1M sodium cacodylate buffer). Brains were placed in a 1 mm brain matrix (Kent Scientific) to generate 1 mm coronal slices that were post-fixed in Karnovsky’s fixative for 2h at ∼21°C, and rinsed with and stored in 0.1 M sodium cacodylate buffer overnight. The medial part of the corpus callosum (∼Bregma +0.5 to -0.5) was dissected and incubated in 0.1 M sodium cacodylate buffer containing 1% (v/v) osmium tetroxide and 1.5% (v/v) potassium ferricyanide [OsO_4_ / K_3_Fe(III)(CN)_6_] in in the dark for 2h at 4°C, before being dehydrated in ethanol and propylene oxide, and embedded in Epon812 resin. Ultrathin 70 nm sections were generated using a Leica Ultra-cut UCT7 and stained with uranyl acetate and lead citrate. Images at 10,000x magnification were collected at 80kV on a Hitachi HT7700 transmission electron microscope. Image collection and analysis was performed by an experimenter blind to the treatment group. Images were opened in Image J (NIH) and axons identified by the presence of microtubules and, for myelinated axons, the presence of an electron dense ring, inner tongue and periaxonal space. The density of axons was quantified from ≥ 10 images per mouse. To quantify axons showing evidence of degeneration, we evaluated ≥ 100 axons per mouse. Axon degeneration was defined by the presence of electron dense patches or clumps of myelin within the axon (termed inclusions) or dark axons (whole axon degeneration), as previously described (Lee *et al*., 2012). Inclusions, by definition, only occur in myelinated axons (Lee *et al*., 2012), but it was not possible to reliably classify whole degenerated axons as myelinated or unmyelinated.

### RNA analyses

To detect *Grik4* mRNA by RNA Scope, brain tissue was collected and cryopreserved as outlined for immunohistochemistry, using solutions pre-treated with 0.1% (v/v) diethyl pyrocarbonate (DEPC; Sigma). 20 µm coronal brain cryosections containing the primary motor cortex or frontal hippocampus (∼Bregma -1.5), and transverse thoracic spinal cord sections (between T1 and T13) were collected as free-floating sections into DEPC-treated PBS. Sections were floated onto RNase-free slides and allowed to dry at ∼21°C for 1 hour. Cryosections were then dehydrated using a 70% to 100% ethanol gradient and dried for 30 min at ∼21°C. *In situ* hybridisation was performed using a RNAscope™ 2.5 HD RED Detection kit according to the manufacturer’s instruction using probes for *Grik4* (In Vitro Technologies, RNAscope® 2.5 Probe - Mm-Grik4-C1; 442021-C1) and a HybEZ hybridisation oven. Slides were washed with DEPC treated milli-Q water and mounted with permafluor mounting medium (Sakura).

Images spanning each brain or spinal cord cryosection were collected using an Axiocam 506 light microscope (Carl Zeiss Microscopy GmbH) with a 2.5 x objective. Adjacent images were manually aligned using Photoshop (Adobe) to generate a single image of the cryosection. Within each cryosection, the anatomical region of interest was defined with reference to anatomical landmarks in the Allen mouse brain atlas. The density of *Grik4* mRNA puncta within each region was determined by performing a particle analysis in Image J (NIH). When puncta were clustered, the approximate number of puncta within a cluster was elucidated by dividing the area of the puncta cluster by the area of a single puncta (0.062 µm^2^, based off the minimum measured size of visually confirmed puncta).

For RNA sequencing, WT and *Grik4^-/-^* mice (P333-P434) were killed by cervical dislocation, their brains removed with chilled dissection instruments and the hippocampus dissected on ice. Tissue was immediately immersed in ice-cold RNAlater™ (Invitrogen, ThermoFisher) and stored at -20°C. Tissue was homogenized in QIAzol lysis reagent using a TissueLyser LT bead beater (QIAGEN), and RNA was extracted using a QIAGEN RNeasy lipid tissue kit (Cat #74804) with on-column DNase I (Cat #79254) treatment following the manufacturer’s instructions. RNA quality was assessed with RNA ScreenTape (Agilent, Cat #5067-5576, 5067-5577 and 5067-5578) using an Agilent 4200 TapeStation. Samples with an RNA integrity number (RIN) >7 were sent to the Australian Genome Research Facility for bulk RNA sequencing. Libraries were generated with the Illumina Stranded mRNA workflow with polyA capture, and samples sequenced to a depth of ≥20M reads with 150 base pair end lengths. The RNA-Seq data, including raw fastq reads, are deposited in NCBI (https://www.ncbi.nlm.nih.gov/) under BioProject ID: PRJNA1076653 (https://www.ncbi.nlm.nih.gov/bioproject/PRJNA1076653). Quality control of raw sequencing data was conducted using Cutadapt (v3.2) (Martin, 2011) within the TrimGalore wrapper (v0.6.7). The Illumina sequencing adapter (5’-CTGTCTCTTATACACATCT - 3’) was trimmed from each read and 1 base pair removed from the 5’ end using Cutadapt. A read quality Phred score threshold of 20 was used to remove low quality reads. Reads < 25 base pairs after trimming were discarded. The quality of the sequencing data was then evaluated using FastQC (v.0.11.9) (Martin, 2011), and all data met the necessary quality metrics for analysis.

Gene expression was quantified using Salmon in mapping-based mode (v.1.6.0) using the Salmon indexed *Mus musculus* (mm10) cDNA reference transcriptome obtained from RefGenie (v0.12.1) (Stolarczyk *et al*., 2020). The sequencing library type was ISR. Within Salmon, --validateMappings was employed to improve the sensitivity and specificity of read mapping, and --seqBias and --gcBias were employed to enable Salmon to learn and correct for sequence-specific biases and fragment-level GC biases respectively.

Differential gene expression analysis was performed using DESeq2 (v1.40.2) (Love *et al*., 2014) to compare gene expression in the hippocampus of WT and *Grik4^-/-^* mice. Tximport (1.28.0) was used to import transcript-level abundance into R (v.4.3.0) and summarise abundance to the gene-level. Genes with < 10 reads across all samples were filtered out. A first pass differential gene expression was performed comparing the *Grik4^-/-^* to the WT samples. Following this, EDASeq (v2.28.0) (Risso *et al*., 2011) was used to account for lane distributional differences in read counts and a threshold p >0.5 determined genes not differentially expressed between groups. These were used as *in silico* empirical negative control genes to calculate 2 factors of unwanted variation using RUVSeq (v1.34.0) (Risso *et al*., 2014), which were included as covariates in the differential gene expression analysis model. Differential gene expression was tested under a hypothesis of log2 fold changes >0.1, with a false discovery rate threshold of alpha = 0.05. Differential gene expression analysis was visualised using EnhancedVolcano (v.1.18.0) (Blighe *et al*., 2023).

### Electrophysiological recordings

#### Compound action potential recordings

Compound action potential (CAP) recordings and conduction-velocity measurements were performed as previously described (Crawford *et al*., 2009; Cullen *et al*., 2021). Briefly, P120-180 mice were euthanized by cervical dislocation and their brains rapidly dissected into ice-cold sucrose solution containing: 75 mM sucrose, 87 mM NaCl, 2.5 mM KCl, 1.25 mM NaH_2_PO_4_, 25 mM NaHCO_3_, 7 mM MgCl_2_, and 0.95 mM CaCl_2_. 400 µm live coronal brain vibratome (Leica VT1200s) slices were generated spanning Bregma +1.0 and -1.0. All slices were incubated at ∼32°C for 45 min in artificial cerebral spinal fluid (ACSF) containing 119 mM NaCl, 1.6 mM KCl, 1 mM NaH_2_PO4, 26.2 mM NaHCO_3_, 1.4 mM MgCl_2_, 2.4 mM CaCl_2_ and 11 mM glucose (300 ± 5 mOsm / kg), before being transferred to ∼21°C ACSF saturated with 95% O_2_/5% CO_2_.

CAPs were evoked by constant current, stimulus-isolated, square wave pulses (200 ms duration, delivered at 0.2 Hz), using a tungsten bipolar matrix stimulating electrode (FHC; MX21AEW), and detected using glass recording electrodes (1-3 MΩ) filled with 3M NaCl. To quantify CAP amplitude, the asymptotic maximum for the short-latency negative peak (myelinated peak; Mye) was first determined by placing the stimulating and recording electrodes 1 mm apart and varying the intensity of stimulus pulses (0.3–4.0 mA) using an external stimulus isolator (ISO-STIM 1D) before recording at 80% maximum stimulation. To enhance the signal-to-noise ratio, all quantitative electrophysiological analyses were conducted on waveforms that were the average of eight successive sweeps, amplified and filtered (10 kHz low pass bessel) using an Axopatch 200B amplifier (Molecular Devices) and digitized at 100 kHz. The conduction velocity of myelinated and unmyelinated axons was estimated by changing the distance between the stimulating and recording electrodes from 0.5 to 2.5 mm, while holding the stimulus intensity constant (80% maximum). The peak latency of myelinated and unmyelinated axons was measured at each distance and graphed relative to the distance separating the electrodes. A linear regression analysis was performed to yield a slope inverse of the velocity. The average velocity for each CAP component was determined for each animal and this value was used for statistical comparisons.

#### Whole cell patch clamp recordings of kainate receptor currents

Acute coronal brain slices (300 µm) were generated from P60-P70 mice and incubated at ∼32°C for 45 min in ACSF before being transferred to ∼21°C ACSF saturated with 95% O_2_/5% CO2, as previously described for the CAP recordings. The rapid desensitisation of kainate receptors was prevented by incubating slices in ∼21°C ACSF containing 50 μM Concanavalin A (ConA; prepared from 10 mg/mL stock solution in 1 M NaCl) for 20 min, immediately preceding recording. ConA treated slices were transferred to a bath and constantly perfused with ACSF (2 mL/min at 21°C). Recording electrodes were prepared from glass capillaries to have a resistance of 3-6 MΩ when filled with an internal solution containing: 125mM Cs-methanesulfonate, 4 mM NaCl, 3 mM KCl, 1 mM MgCl_2_, 8 mM HEPES, 9 mM EGTA, 10 mM Phosphocreatine, 5 mM MgATP and 1 mM Na_2_GTP and set to a pH of 7.2 with CsOH, and an osmolarity of 290 ± 5 mOsm/kg. Whole cell patch clamp recordings were collected from layer V pyramidal neurons in the motor cortex (pyramidal soma, capacitance > 75 pF, and membrane resistance < 250 mΩ; (Oswald et al. 2013; Suter et al. 2013)), using a HEKA patchclamp EPC800 amplifier and pclamp 10.5 software (Molecular devices). A gap free protocol (sampled at 50 kHz and filtered at 3 kHz) with a holding potential of -70 mV was used to record the effect of kainate. The perfusate was first changed to a solution containing GYKI 52466 (50 μM) and picrotoxin (100 μM, Sigma) to block AMPA and GABA_A_ receptors, respectively. For some recordings, GYKI52466 was exchanged for CNQX (50 μM), which blocks both AMPA and kainate receptors. After 2 min, 50 μM kainate was included in the perfusate for 2 min to activate kainate receptors, before all drugs were washed out. The maximum amplitude of the inwards current evoked by drug application was calculated to determine the kainate-evoked current.

#### Spontaneous and miniature excitatory post synaptic currents

Acute brain slices were prepared as described for CAP recordings, transferred to a bath constantly perfused with ACSF (2 mL/min at 21°C), and whole cell patch clamp recordings obtained from layer V pyramidal neurons at a holding potential of -70 mV. After obtaining whole cell configuration, the perfusate was switched to ACSF containing 100 µM picrotoxin. Spontaneous excitatory post synaptic currents (sEPSC) were recorded after 2 min of perfusion using a 3-minute gap free protocol sampled at 50 Hz and filtered at 1 Hz. Cells were then washed with ACSF containing 100 µM picrotoxin and 500 nM tetrodotoxin (TTX; Abcam) and miniature excitatory post synaptic currents (mEPSC) recorded after 2 min perfusion using a 3 min gap free protocol, sampled at 50 Hz and filtered at 1 Hz. Access resistance was recorded before and after each protocol and recordings were excluded if the access resistance exceeded 20 MΩ. Measurements were made from data files using clampfit 10.5 (molecular devices). sEPSC and mEPSC had an amplitude ≥8 pA and were analysed using the MiniAnalysis60 program (Synaptosoft, Decateur, Georgia). Post-recording, the patch electrode was removed and pyramidal neuron identity and location within layer V of the motor cortex was visually identified following detection of neurobiotin with a fluorescently conjugated Alexa Fluor® 546 streptavidin (Molecular Probes™; 0.1% triton-X-diluent detergent).

#### Action potential measurements

Acute coronal brain slices were generated from P120-180 mice and transferred to a bath constantly perfused with ACSF (2 mL/min; ∼21°C). Recording electrodes were prepared from glass capillaries to have a resistance of 3-6 MΩ when filled with an internal solution containing 130 mM K-gluconate, 4 mM NaCl, 10 mM HEPES, 0.5 mM CaCl_2_, 10 mM BAPTA, 4 mM MgATP, 0.5 mM Na_2_GTP, pH set to 7.4 with KOH and an osmolarity of 290 ± 5 mOsm/kg. Whole cell patch clamp recordings of layer V motor cortex pyramidal neurons were collected using a HEKA patchclamp EPC800 amplifier and pclamp 10.5 software (Molecular devices). Current was applied in current clamp configuration so that the cells rested at -70 mV, and a series of current steps were applied in 20 pA increments (0 to 260 pA) to evoke action potentials and determine the firing patterns of the cell. Following this, current was injected so that cells were resting at -90 mV and a single action potential, evoked by a square 300 pA pulse, recorded. To enhance the signal-to-noise ratio, the quantification was performed using the average of eight successive sweeps, filtered at 3 kHz. Action potential parameters were analysed and quantified using Clampfit 11.1 software (Molecular devices). Neuron firing patterns were obtained by counting the number of action potentials fired at each current step. Action potential threshold was calculated using the trace derivative to find the point where the change in voltage reaches 20 mV/ms, reflecting rapid sodium channel opening; initiation time was measured as the time between initiation of the current injection and threshold; rheobase was obtained by measuring the current necessary to pass the single action potential threshold; rise time was calculated as the time taken from threshold to reach maximum action potential amplitude; amplitude was taken from the action potential threshold to the peak; area under the curve (AUC) was obtained by calculating the integral of the peak amplitude using the software; half-width was derived by measuring the time when rising or decaying slopes are at 50% of their maximum; decay-time was measured as the time elapsed between peak amplitude and a return to baseline; the after hyperpolarization was measured as the voltage difference between the peak hyperpolarization and baseline. The baseline was defined as the voltage recorded prior to application of the depolarising current step.

### Statistical analyses

All statistical analyses were performed using Prism version 8 or 10 (GraphPad Software). The Shapiro-Wilk test was used to determine if the datasets followed a normal distribution. Data comparing two groups at a single time point were analysed using a parametric two-tailed t-test or a non-parametric Mann-Whitney U test. For data with more than two comparisons, 1- or 2-way ANOVAs with or without repeated measures were used as appropriate, with either Bonferroni or Šídák multiple comparison posthoc tests. Where data are presented as cumulative distribution plots, data were analysed using a Kolmogorov–Smirnov (KS) test. Change in CAP amplitude with increasing stimulus strength (mV) was analysed by linear regression. As DigiGait treadmill data were not available for all mice at every timepoint (unwillingness to run became more prevalent with age), gait parameters were analysed using a restricted maximum likelihood (REML) mixed effects model with a Geisser-Greenhouse correction, followed by a Bonferroni post-test. Data from the beam walk and grip strength tests were also analysed using a Geisser-Greenhouse correction, as to not assume equal variability and sphericity. A Grubbs’ test was used to remove significant (p<0.05) outliers. This resulted in the removal of one WT mouse from our analysis of somatic presynaptic sites. Details relevant to each statistical analysis, including the number of mice analysed in each group, or the number of cells, axons, or puncta quantified, is provided in the corresponding figure legend.

### Key resources table

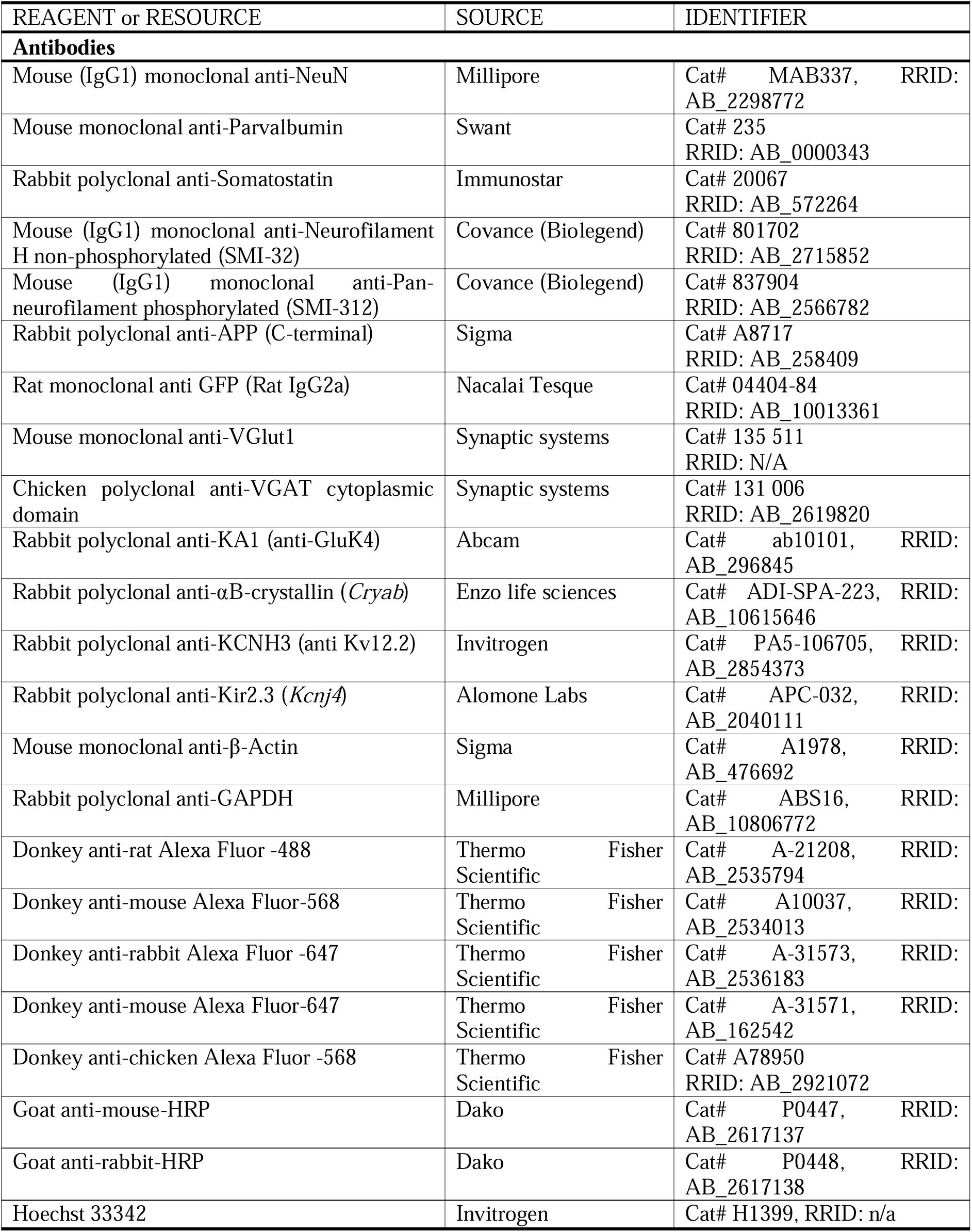
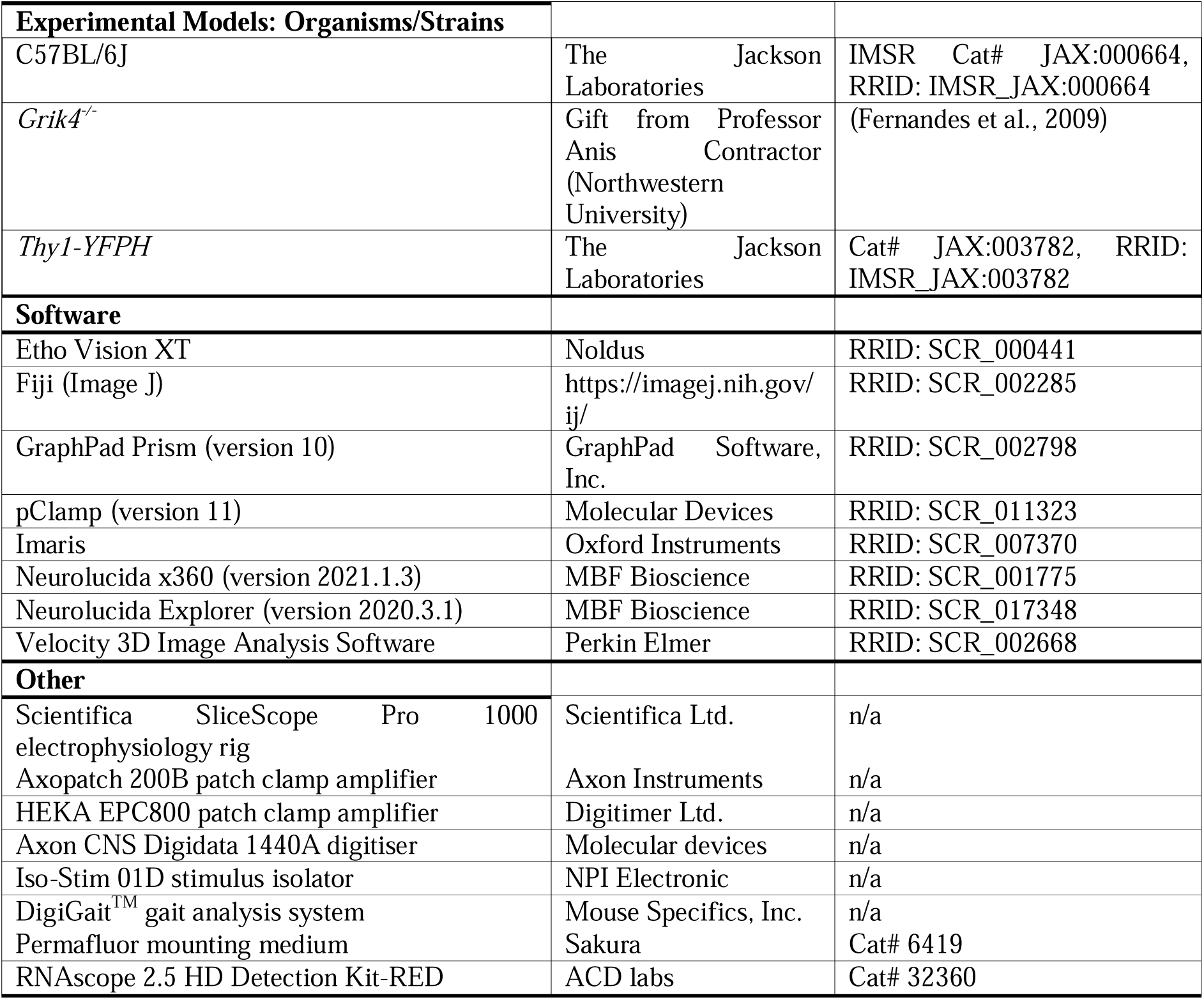

## Results

### Motor cortex layer V pyramidal neurons express functional Gluk4-containing kainate receptors

Gluk4-containing kainate receptors are highly expressed in the hippocampus (Friedman *et al*., 1994; Wisden and Seeburg, 1993) and while some evidence supports a wider expression pattern (Arora *et al*., 2018), the level of *Grik4* expression in the cortex has been controversial (Arora *et al*., 2018; Hadzic *et al*., 2017; Kask *et al*., 2000; Lein *et al*., 2007). To quantify the expression of *Grik4* mRNA in regions of the cortico-motor circuit, we performed RNA Scope *in situ* hybridisation on 20 µm coronal brain and transverse thoracic spinal cord cryosections from P180 adult mice **(Figure 1 a-d)**. Discrete *Grik4* mRNA puncta (red) were detected in the hippocampus (**Figure 1e**), which served as our experimental positive control, with a particularly high density of puncta detected over the neuronal cell body layer of the CA3 (**Figure 1a, f**). The spinal cord also contained cells, primarily within grey matter, that expressed *Grik4* (**Figure 1b, e**), including large cells within the ventral horn, presumably lower motor neurons, and cells within the dorsal horn – an area which contains interneurons that integrate sensory information and relays this to the motor neurons (**Figure 1b**). *Grik4* puncta were also expressed by cells throughout the adult mouse motor cortex (**Figure 1d, e**) including cells within layers II/III, V and VI, suggesting localisation with the pyramidal neurons of the motor cortex (**Figure 1d, g**).

**Figure 1.**
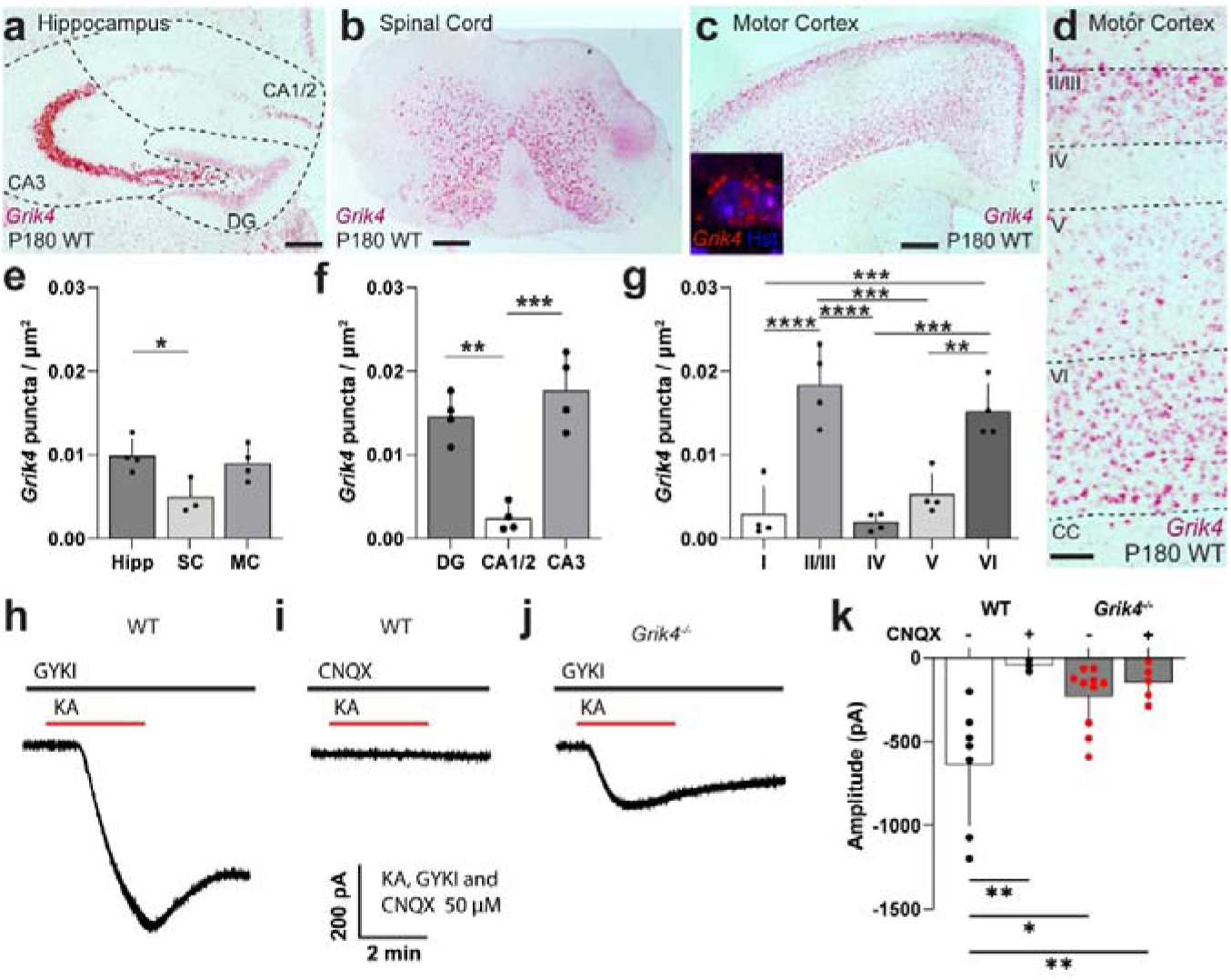
*Grik4* is expressed in CNS motor regions. **a-d)** RNA Scope *in situ* hybridisation showing *Grik4* (red) expression in the hippocampus (**a**, positive control), spinal cord (**b**) and motor cortex (**c**, inset shows high magnification image of a Hoechst^+^ nuclei with *Grik4^+^* puncta; **d**, motor cortex layers). **e)** Average number of *Grik4* puncta per μm^2^ in the hippocampus, spinal cord and motor cortex [One-way ANOVA, F(2,8) = 5.500; p=0.031]. **f)** Average number of *Grik4* puncta per μm^2^ in the dentate gyrus, CA1/2, or CA3 [One-way ANOVA, F(2,9) = 25.68; p<0.0002]. **g)** Average number of *Grik4* puncta per μm^2^ in motor cortex layers I-VI [One-way ANOVA, F(4,15) = 21.94; p<0.0001]. **h-j)** Whole cell patch clamp current traces from pyramidal neurons in layer V of the motor cortex of WT or *Grik4^-/-^* mice as kainate (50 μM) is applied in the presence of GYKI (AMPA receptor blocker) or CNQX (AMPA and kainate receptor blocker). **k)** Average amplitude of the kainate-induced current in layer V pyramidal neurons in the motor cortex of WT or *Grik4^-/-^* mice, evoked in the presence (+) or absence (-) of CNQX [2-way ANOVA: genotype F(1,21) = 2.169, p = 0.1557; antagonist F(1,21) = 10.60, p = 0.0038; interaction F(1,21) = 6.066, p = 0.0225]. Data are presented as mean ± SD. Bonferroni’s or Tukey’s multiple comparisons post-test results shown as * p = < 0.05, ** p = < 0.01, *** p = < 0.001 and **** p = < 0.0001. n=3-4 mice per genotype (**e-g**) or 3-10 cells from 3-4 mice per genotype (**k**). Scale bars represent: 0.3mm (a, c) or 0.1mm (b, d).

To determine whether the expression of *Grik4* mRNA puncta, even at moderate levels, is associated with the formation of functional GluK4-containing kainate receptors, we performed whole cell patch clamp electrophysiology to compare kainite-receptor mediated currents in layer V pyramidal neurons in the motor cortex of WT and GluK4 functional knockout mice (*Grik4^-/-^*) (Fernandes *et al*., 2009). *Grik4^-/-^* mice lack exon 16 of the *Grik4* gene (validated by PCR and Sanger sequencing) but are a normal body weight and are fertile (**Figure S1**). In these mice, Gluk4 is expressed at WT levels in brain protein lysates, and the loss of the pore-forming sequence does not change protein size enough to be notable by Western blot (**Figure S1**). However, the resulting Gluk4-containing kainate receptors are not able to conduct ions (Fernandes *et al*., 2009). Currents were evoked by bath applying kainate to ConA treated brain slices in the presence of GYKI 52466, which selectively blocks AMPA receptor activation (**Figure 1h-j)**. In WT mice, this resulted in an inward current that could be blocked by the AMPA/kainate receptor antagonist CNQX (**Figure 1h, i, k**). A kainate-specific current was still evoked in *Grik4^-/-^* mice but was reduced in amplitude by ∼64% (**Figure 1j-k**). These data indicate that GluK4-containing kainate receptors are functionally expressed by layer V pyramidal neurons in the motor cortex, and account for the majority of their kainate receptor mediated current.

### Grik4^-/-^ mice are hyperactive and have impaired motor coordination

*Grik4^-/-^* mice have a reduction in depressive- and anxiety-like behaviours in the forced swim test and elevated plus maze (Catches *et al*., 2012), however, the outcome of these tests is heavily reliant on motor performance. As the loss of Gluk4-containing kainate receptors from the motor cortex and other levels of the cortico-motor circuit could impact the outcome of these tests, we subjected adult WT and *Grik4*^-/-^ mice to a battery of behavioural tests, to evaluate affective but also locomotor behaviour (**Figure 2a**). In the Y-maze, P60 WT and *Grik4*^-/-^ mice (n ≥9 mice per group) explored the maze, and spent more time in the novel arm, relative to the initial or second arms (**Figure 2d**), indicating that short term memory is normal. However, *Grik4^-/-^*mice had an increased number of Y-maze arm entries compared to WT mice, which reached significance for the novel arm (**Figure 2e**), suggestive of an increased level of activity. In the elevated plus maze, *Grik4*^-/-^ mice spent more time in the open maze arms than WT mice, suggesting they were less prone to anxiety-like behaviour (**Figure 2f-h**). However, when tested with the marble bury test, which is designed to evaluate anxiety and obsessive or compulsive behaviours, *Grik4^-/-^* and WT mice buried an equivalent number of marbles (**Figure 2j**). As with the Y-maze task, in the elevated plus maze *Grik4*^-/-^ mice entered the maze arms more frequently than WT mice (**Figure 2i**), suggesting that part of this phenotype may reflect hyperactivity. When subjected to a forced swim, *Grik4*^-/-^ mice spent less time immobile (**Figure 2j**) and more time moving before their first period of immobility (**Figure 2k**), suggesting that they are less prone to depressive-like behaviour, but again, this may be influenced by a generalised increased in activity.

**Figure 2.**
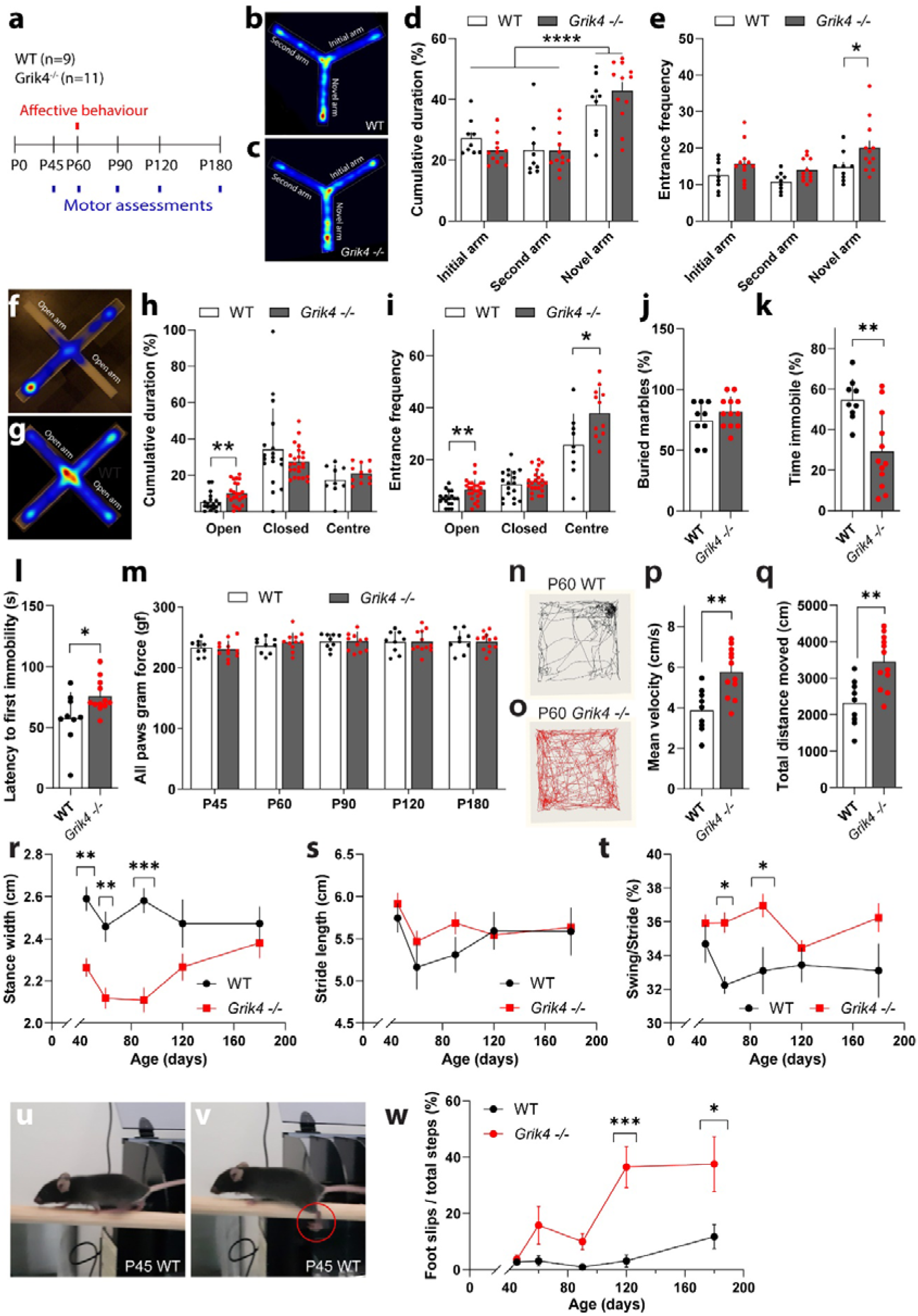
Gluk4 knock-out mice are hyperactive and have impaired motor coordination. **a)** Timeline for affective and motor behavioural testing. **b-c)** Heat maps generated from a WT (b) and *Grik4^-/-^* (c) mouse showing the relative time spent in each area of the Y-maze. **d)** The proportion of time (% of total) that WT and *Grik4^-/-^* mice spent in the initial, second or novel arm of the Y-maze [2- way ANOVA: Time F(2,57) = 29.43, p<0.0001; Genotype F(1,57) = 0.0053, p=0.9417; Interaction F(2,57) = 1.614, p=0.2080]. **e)** The number of times that WT or *Grik4^-/-^* mice entered the initial, second or novel arm of the Y maze [2-way ANOVA: Frequency F(2,57) = 6.068, p<0.0041; Genotype F(1,57) = 10.17, p<0.0023; Interaction F(2,57) = 0.3398, p = 0.71]. **f, g)** Heat maps showing the relative time that the WT (**f**) or *Grik4^-/-^*(**g**) mouse spent in each region of the elevated plus maze. **h)** The proportion of time (% of total) that WT and *Grik4^-/-^* mice spent in the open arms, closed arms or centre of the elevated plus maze [2-way ANOVA: Time F(2,99) = 42.99, p<0.0001; Genotype F(1,99) = 0.047, p=0.83; Interaction F(2,99) = 3.005, p = 0.054]. **i)** The number of times that WT or *Grik4^-/-^* mice entered the open arms, closed arms or central region of the elevated plus maze [2-way ANOVA: Frequency F(2,99) = 121.5, p<0.0001; Genotype F(1,99) = 19.76, p<0.0001; Interaction F(2,99) = 5.761, p = 0.004]. **j)** The proportion of the marble that were buried by WT and *Grik4^-/-^* mice [unpaired t-test, t = 1.16, p = 0.26]. **k)** The proportion of time (% of total) that WT and *Grik4^-/-^* mice spent immobile during the forced swim test [unpaired t-test, t = 3.627, p = 0.0018]. **l)** The latency between commencing the forced swim test and the first period of immobility for WT and *Grik4^-/-^* mice [unpaired t-test, t = 2.351, p = 0.0296]. **m)** Average force required at which WT or *Grik4^-/-^*mice release the bar in the grip strength test [Repeated measures 2-way ANOVA: Age F(3.3,62.75) = 2.443, p=0.067; Genotype F(1,19) = 0.0253, p = 0.975; Interaction F(4,74) = 0.2409, p = 0.91]. **n, o)** Map of the path taken by a WT (**n**) and *Grik4^-/-^* (**o**) mouse exploring the open field. **p)** The mean velocity of WT and *Grik4^-/-^*mice exploring the open field [unpaired t-test, t = 3.745, p=0.0014]. **q)** The total distance travelled by WT and *Grik4^-/-^* mice exploring the open field [unpaired t-test, t=3.743, p=0.0014]. **r)** The average stance width of WT and *Grik4^-/-^* mice running on the Digigait at 22cm/sec [2-way ANOVA: Age F(4,77) = 1.532, p=0.2012; Genotype F(1,77) = 43.13, p<0.0001; Interaction F(4,77) = 2.053, p=0.095]. **s)** The average stride length of WT and *Grik4^-/-^*mice running on the Digigait at 22cm/sec [2-way ANOVA: Age F(4,77) = 2.313, p=0.065; Genotype F(1,77) = 2.164, p=0.144; Interaction F(4,77) = 0.431, p=0.786]. **t)** Swing time expressed as a percentage of total stride time for WT and *Grik4^-/-^*mice running on the Digigait at 22cm/sec [2-way ANOVA: Age F(4,77) = 0.919, p=0.457; Genotype F(1,77) = 21.53, p<0.0001; Interaction F(4,77) = 1.214, p=0.312]. **u, v)** Image of a P45 WT (**u**) or *Grik4^-/-^*(**v**) mouse crossing the beam. The red circle denotes a foot slip, where the foot goes below the level of the beam. **w)** The proportion (%) of steps taken by WT or *Grik4^-/-^* mice that result in a foot slip in the beam crossing task [2-way ANOVA: Age F(4,93) = 5.453, p=0.0005; Genotype F(1,93) = 21.69, p<0.0001; Interaction F(4,93) = 2.808, p=0.03]. Data are presented as mean ± SD (bar graphs) or mean ± SEM (line graphs). Bonferroni post-test or t-test outcomes are presented as: * p<0.05, ** p<0.01, *** p<0.001 or **** p<0.0001. Data are from n=9 WT and n=11 *Grik4^-/-^*mice.

As *Grik4* mRNA was highly expressed in the motor cortex and spinal ventral horn, we next aimed to determine whether Gluk4-containing kainite receptors are critical for normal motor function between P45 and P180 (**Figure 2a**). Across this time course, the grip strength of all four paws was unaffected by genotype (**Figure 2m**), indicating that muscle strength is not regulated by Gluk4-containing kainite receptors. At P60, when WT and *Grik4^-/-^* mice were placed in an open field arena (**Figure 2n, o**) and their free movement tracked, the locomotor activity of *Grik4^-/-^* mice was elevated, as the mice moved more quickly and covered a greater distance than WT mice (**Figure 2p**, q). To directly compare the gait of mice in each group, we controlled their speed of movement - placing WT and *Grik4^-/-^* mice on the Digigait treadmill, with the belt speed set to 22 cm/s. We measured forelimb and hindlimb gait parameters and expressed each as the average of the left and right paws. While forelimb gait appeared normal (**Figure S2**), hindlimb movement was significantly altered in the *Grik4^-/-^* mice (**Figure 2r-t)**. Hindlimb stance width was reduced (**Figure 2r),** stride length was unchanged (**Figure 2s**), but swing time formed a higher proportion of the stride (**Figure 2t**), a change typically observed when moving at a greater speed (Hebenstreit *et al*., 2015). This phenotype was most striking prior to 4 months of age, as ∼30% of mice fail to run on the treadmill and cannot be analysed ≥P120. Indeed, it is unlikely that the gait phenotype detected in *Grik4^-/-^*mice reduced with ageing, as their performance in the beam walk test became increasingly worse with age (**Figure 2u-w**). At P45, WT and *Grik4*^-/-^ mice recorded an equivalent number of foot slips as they walked across the suspended beam, however, at P120 and P180, *Grik4^-/-^* mice made approximately twice as many foot slips as WT mice (**Figure 2w**), indicating that their balance and/or fine motor coordination was increasingly impaired.

### Action potential properties are normal in layer V pyramidal neurons of Grik4^-/-^ mice

To determine how the loss of functional GluK4-containing kainate receptors influences neuronal function within the motor circuit, at an age when motor performance was impaired, we performed whole cell patch current clamp analysis of layer V pyramidal neurons in the motor cortex of P180 mice. The firing threshold and frequency of action potential firing in response to current steps were equivalent between WT and *Grik4^-/-^* mice (**Figure 3a-c**). Furthermore, when a single action potential was evoked (**Figure 3d, e**), the properties of that action potential, including: action potential initiation time (**Figure 3f**), rheobase (**Figure 3g**), threshold (**Figure 3h**), rise time (**Figure 3i**), amplitude (**Figure 3j**), area under the action potential curve (AUC; **Figure 3k**), half-width (**Figure 3l**), decay time (**Figure 3m**) and after hyperpolarisation amplitude (**Figure 3n**) were unchanged, indicating that layer V motor cortex neurons in *Grik4^-/-^* mice elicit normal action potentials.

**Figure 3.**
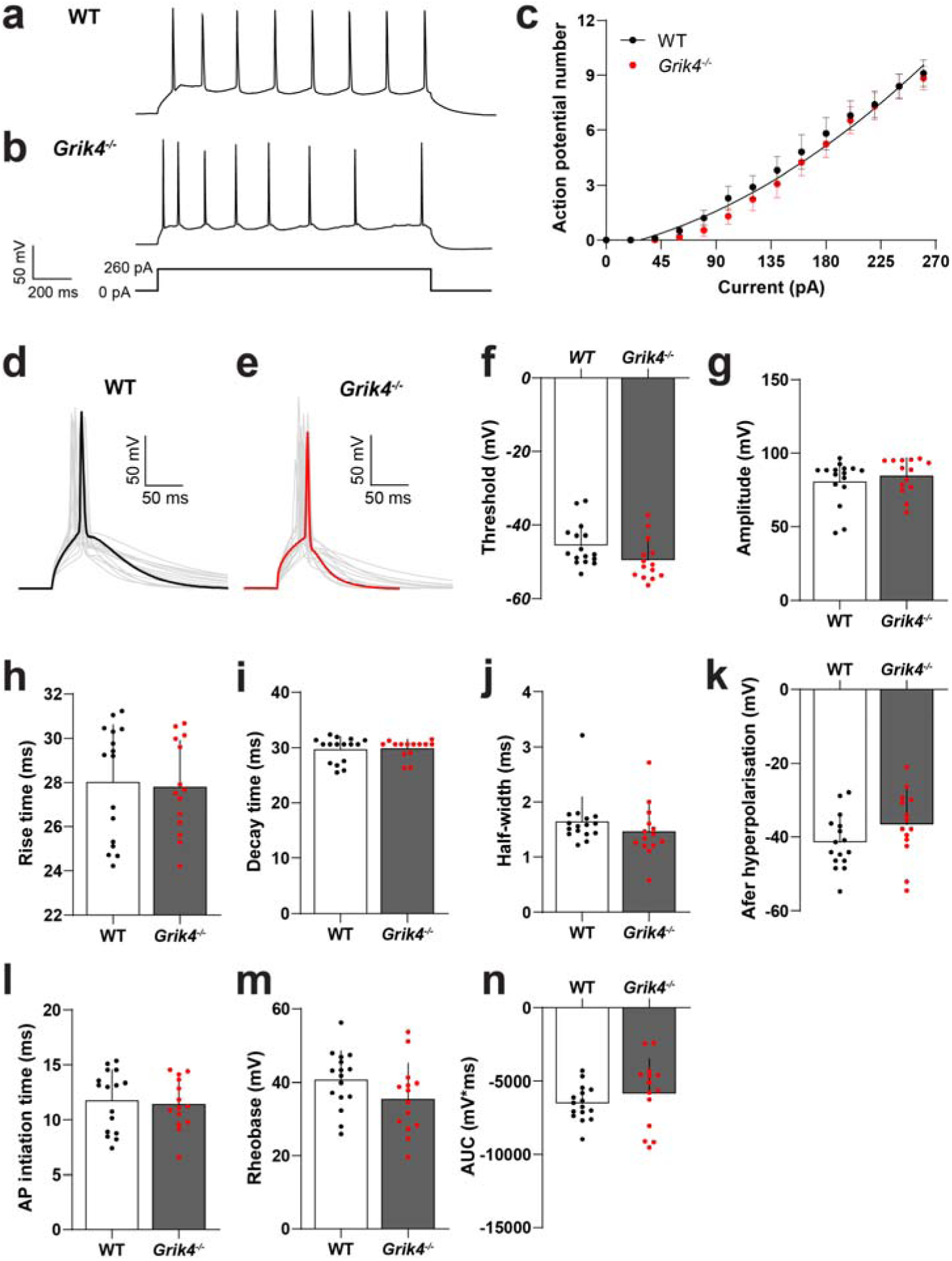
The firing properties of layer V motor cortex pyramidal neurons is normal in *Grik4^-/-^* mice. **a-b)** Example traces of action potentials from layer V pyramidal neurons in the motor cortex of WT (**a**) or *Grik4^-/-^*(**b**) mice, evoked in response to a 1s and 260 pA current step. **c)** The number of action potentials evoked in layer V pyramidal neurons in WT or *Grik4^-/-^* in response to current steps [curve fit, F(3, 316) = 1.424, p=0.236, r^2^=0.7248]. **d-e)** Voltage traces (grey) showing action potentials evoked in WT (**d**, black) and *Grik4^-/-^* (**e**, red) layer V pyramidal neurons by a 100 ms and 300 pA current step. **f)** Action potential threshold [t-test, t=1.923, p=0.065]. **g)** Action potential amplitude [t-test, t=0.8031, p=0.43]. **h)** Action potential rise time [t-test, t=0.2563, p=0.8]. **i)** Action potential decay time [t-test, t=0.2406, p=0.8116]. **j)** Action potential half-width [t-test, t=1.061, p=0.298]. **k)** Action potential afterhyperpolarisation amplitude [t-test, t=1.597, p=0.122]. **l)** Action potential initiation time [t-test, t=0.3463, p=0.732]. **m)** Rheobase [t-test, t=1.619, p=0.117]. **n)** Area under the curve (AUC) of the action potential [t-test, t=1.021, p=0.316]. Data presented as mean ± SEM (c) or mean ± SD (f-n) for n=10-16 cells from 4-5 mice in each group.

### Grik4^-/-^ mice have fewer excitatory synapses onto layer V pyramidal neurons in the motor cortex

As the excitability of the pyramidal neurons appeared normal in *Grik4*^-/-^ mice, we next examined the inputs onto these neurons. In the layer V pyramidal neurons of P60 WT and *Grik4^-/-^* mice, we recorded mEPSCs in the presence of tetrodotoxin to block action potentials, and sEPSCs, under conditions where neurons can fire action potentials (**Figure 4a-b**). The frequency of mEPSCs received by layer V pyramidal neurons was reduced by ∼50% in *Grik4^-/-^* compared with WT mice (**Figure 4c**). sEPSC frequency was also reduced by a similar magnitude (**Figure 4c**). Furthermore, there was no change in the amplitude of the mEPSCs or sEPSCs (**Figure 4d**), however, the decay time of both mEPSCs and sEPSCs was increased in the *Grik4*^-/-^ mice (**Figure 4e-h**). As the kinetics of kainate receptor-mediated EPSCs are typically slower than the kinetics of other glutamate receptors (Castillo *et al*., 1997; Kidd and Isaac, 2001), it is unlikely that this increase in decay time results directly from losing a GluK4-containing kainate receptor component of the EPSC. Instead, it may reflect an altered excitatory receptor subunit profile at the remaining synapses.

**Figure 4.**
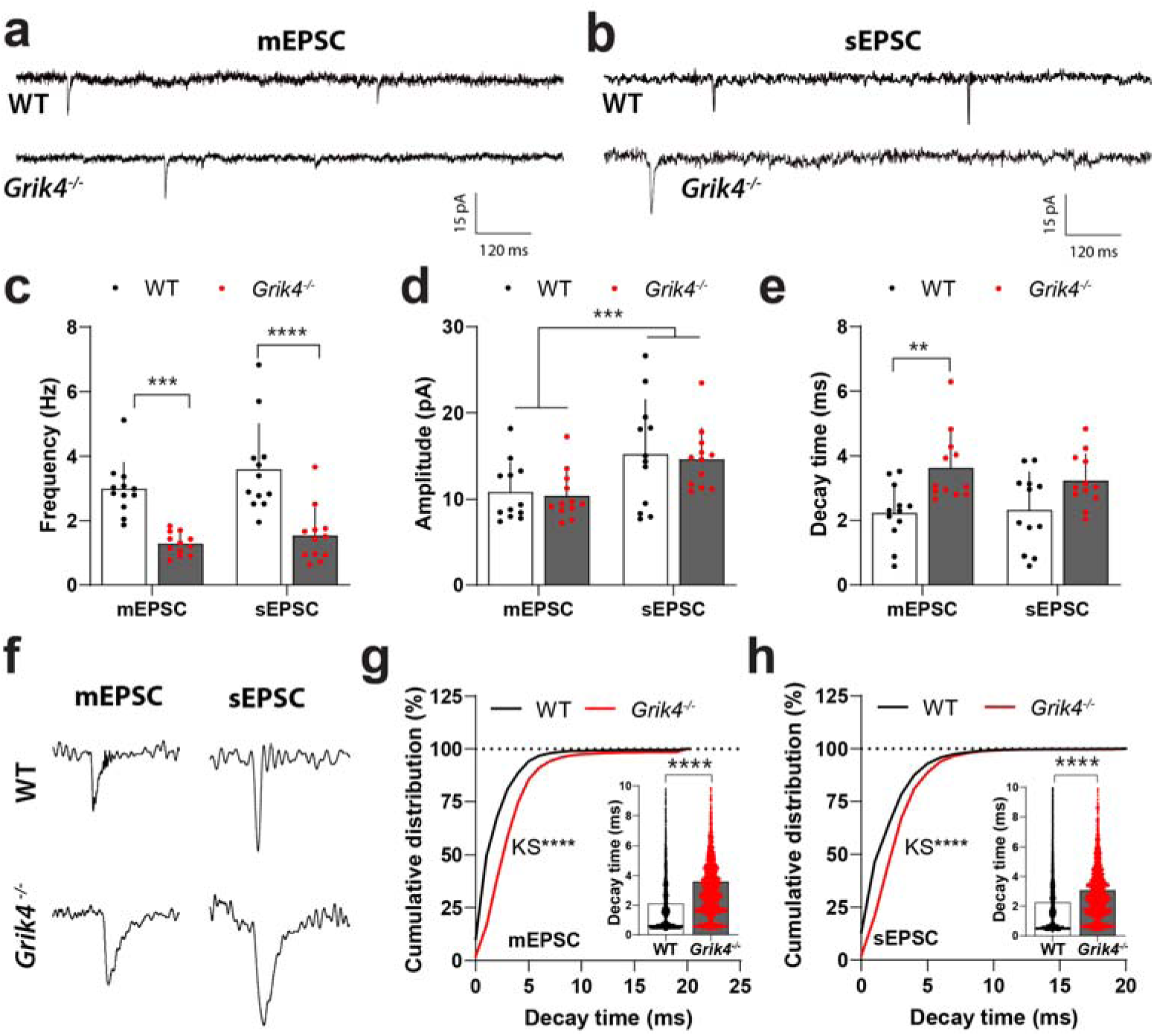
Excitatory post-synaptic currents are less frequent in layer V pyramidal neurons of Grik4^-/-^ mice. **a)** Current traces showing miniature excitatory post-synaptic currents (mEPSCs), recorded in the presence of TTX, from layer V pyramidal neurons of WT and *Grik4^-/-^* mice. **b)** Current traces showing spontaneous excitatory post-synaptic currents (sEPSCs) recorded from layer V pyramidal neurons of WT and *Grik4^-/-^*mice. **c)** The frequency of excitatory events recorded from layer V pyramidal neurons in WT and *Grik4^-/-^*mice [2-way ANOVA: Event type F(1,44) = 2.525, p=0.119; Genotype F(1,44) = 47.90, p<0.0001; Interaction F(1,44) = 0.4075, p=0.527]. **d)** The mean amplitude of excitatory events recorded in WT and *Grik4^-/-^* neurons [2 way ANOVA: Event type F(1,44) = 12.64, p=0.009; Genotype F(1,44) = 0.1802, p=0.673; Interaction F(1,44) = 0.0036, p=0.95]. **e)** Average mEPSC decay time per cell for neurons in WT and *Grik4^-/-^* mice [2 way ANOVA: Event type F(1,44) = 0.2972, p=0.59; Genotype F(1,44) = 15.75, p=0.0003; Interaction F(1,44) = 0.695, p=0.409]. **f)** Example traces showing individual excitatory events from WT and *Grik4^-/-^* neurons. **g)** Cumulative distribution (scatter plot inset) of all mEPSCs recorded from WT (5819 events) or *Grik4^-/-^* neurons (2761 events) [Kolmogorov-Smirnov D=0.3336, p<0.0001; Mann- Whitney U=4707957, p<0.0001]. **h)** Cumulative distribution (scatter plot inset) of all sEPSCs recorded from WT (7476 events) or *Grik4^-/-^* neurons (3318 events) [Kolmogorov-Smirnov D=0.2592, p<0.0001; Mann-Whitney U=8652696, p<0.0001]. Data presented as mean ± SD. Outcomes of Kolmogorov-Smirnov test, Mann-Whitney or Bonferroni post-test denoted as: ** p < 0.01, *** p < 0.001 or **** p < 0.0001 for recordings made from n=12 cells from 4-6 mice per group.

The change in EPSC frequency detected in *Grik4^-/-^* mice may reflect fewer excitatory synaptic connections onto pyramidal neurons in the motor cortex of *Grik4^-/-^* mice, or a reduction in glutamate release probability at synapses. The overexpression of *Grik4* increases the probability of glutamate release in the amygdala of P17-21 mice, as the increased affinity of presynaptic kainate receptors resulted in their activation by ambient glutamate levels (Arora *et al*., 2018). To distinguish between these possibilities, we quantified synapse number along YFP-labelled basal dendrites elaborated by layer V pyramidal neurons in the motor cortex of control (*Thy1-YFPH*) or *Grik4^-/-^*(*Thy1-YFPH :: Grik4 ^-/-^*) mice. ≥10 μm dendrite segments were imaged and reconstructed, and each spine manually traced/defined (**Fig. 5a-b**), so that spine density and morphology could be analysed. At P60, the density of spines was significantly reduced on the basal dendrites of *Grik4*^-/-^ mice, compared to control mice (**Figure 5c)**. By classifying each dendritic spine as having a thin, stubby or mushroom morphology, we found that *Grik4^-/-^*mice had a significantly reduced density of thin spines (**Figure 5c)** and the proportion of spines that were thin was also reduced (**Figure 5d)**. This was accompanied by a smaller increase in the density of spines classified as mushroom spines (**Figure 5c-d).** Thin spines are relatively dynamic and highly plastic (Pchitskaya and Bezprozvanny, 2020) while mushroom spines are largely stable overtime, having previously undergone synaptic strengthening (Albarran *et al*., 2021; Basu *et al*., 2018). The loss of thin spines may suggest that GluK4-kainate receptors are involved in synapse formation or keeping spines labile. The increase in mushroom spines may represent a homeostatic adjustment of synaptic weights, to allow the smaller number of synapses to respond more strong, and compensate for the overall loss of excitatory input (Fauth and Tetzlaff, 2016; Letellier *et al*., 2019).

**Figure 5.**
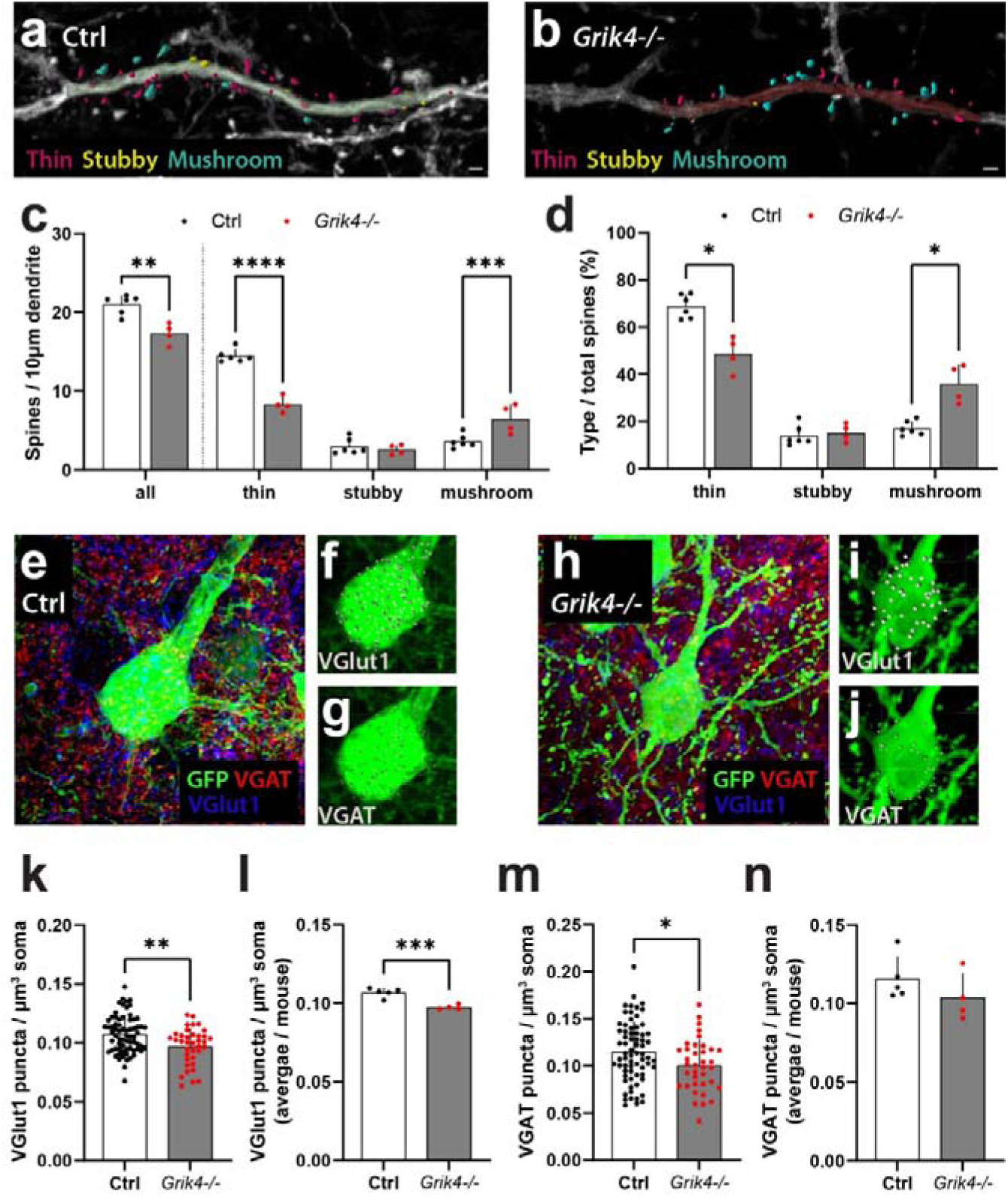
*Grik4^-/-^*layer v pyramidal neurons have a reduced density of basal dendritic spines and somatic excitatory presynaptic contacts. **a-b)** Reconstructed basal dendrite segment from a control (Ctrl, **a**) or *Grik4^-/-^* (**b**) mouse, with spines classified as thin (red), stubby (yellow), or mushroom (blue) morphological subtypes. Scale bars represent 1µm. **c)** Number of spines per 10µm stretch of basal dendrite, average per mouse [unpaired t-test, t = 4.64, p = 0.002]. Number of thin, stubby or mushroom spines per 10µm stretch of basal dendrite, average per mouse [2-way mixed ANOVA: Spine type (within-subjects) F(2,16) = 136.1, p<0.0001; Genotype F(1,8) = 21.50, p = 0.0017; Interaction F(2,16)=35.54, p<0.0001]. **d)** Basal dendritic spine morphological types expressed as percentage of total spines within each dendrite, averaged per mouse [Mann-Whitney U compare mean ranks between genotypes, within each spine type, Bonferroni adjusted p-values; thin rank diff = 5, p=0.0286; stubby rank diff = -1.25, p >0.999; mushroom rank diff = -5, p = 0.029]. e-j) *Thy1-YFPH^+^* neuronal soma in layer V of the motor cortex of a WT (**e-g**) or *Grik4^-/-^* (**h-j**) mouse, stained to detect GFP (green), vesicular GABA Transporter (VGAT; red; marking inhibitory synaptic terminals), and vesicular glutamate transporter 1 (VGlut1; blue; marking excitatory synaptic terminals). Imaris software was used to identify and mark puncta that were close to the GFP^+^ neuronal surface (grey spots in **f**, **g**, **i** and **j**). **k)** The density of VGlut1 puncta in GFP^+^ Ctrl neurons (n=69) and *Grik4^-/-^* neurons (n=37) [Unpaired t-test, t=3.281, P=0.0014]. **l)** The density of VGlut1 puncta at GFP^+^ neuron soma, average per mouse [Unpaired t-test t=5.760, p=0.0007]. **m)** The density of VGAT puncta in individual GFP^+^ neurons of Ctrl mice (n=69 cells) and *Grik4^-/-^* mice (n=37 cells) [Unpaired t-test, t = 2.342, p = 0.021]. **n)** The density of VGAT puncta at GFP^+^ neuron soma, average per mouse [Unpaired t-test t=1.196, p=0.2707]. Data presented as mean ± SD. Outcome of Šídák’s multiple comparisons tests (c), Bonferroni-corrected Mann-Whitney U tests (d) or t-tests (k-n): * p<0.05, **p<0.01, *** p<0.001 or **** p<0.0001. n = 4-6 mice in each group.

The loss of excitatory dendritic spines was also mirrored by a loss of excitatory presynaptic terminals on the soma of layer V pyramidal neurons in the motor cortex of *Grik4 ^-/-^* mice. By performing immunohistochemistry on P60 control (*Thy1-YFPH*) and *Grik4^-/-^*(*Thy1-YFPH :: Grik4 ^-/-^*) mice to detect the expression of a vesicular transporter for glutamate, VGlut1, we were able to label and quantify the number of excitatory presynaptic terminals in close proximity to individual YFP^+^ pyramidal neuronal soma (**Figure 5e-f, h-i**). The density of VGlut1^+^ puncta proximal to individual YFP^+^ soma was significantly reduced in *Grik4^-/-^*mice (**Figure 5k**), as was the average density of VGlut1^+^ puncta per YFP^+^ soma per mouse (**Figure 5l**), suggesting that fewer excitatory presynaptic terminals release glutamate onto the soma of these neurons. As neuronal output is regulated by excitatory and inhibitory input, we also examined the expression a vesicular transporter of GABA (VGAT; **Figure 5 e, g, h, j**). The density of VGAT^+^ puncta proximal to the YFP^+^ soma of layer V pyramidal neurons, sampled from *Grik4^-/-^* mice, suggested a reduction in the density of inhibitory presynaptic sites (**Figure 5m**), however, the biological effect was less robust than the loss of excitatory terminals, as the average density of VGAT^+^ puncta per YFP^+^ soma per mouse was unaffected by genotype (**Figure 5n)**.

### Grik4^-/-^ mice have a lower callosal axon density at P180 but do not show signs of overt axon degeneration

To determine whether a reduction in excitatory synapse number was the result of *Grik4^-/-^* mice having fewer neurons in the motor cortex, coronal brain cryosections from P20, P60 and P180 WT and *Grik4^-/-^* mice were processed to detect NeuN, parvalbumin and somatostatin. However, the loss of functional Gluk4-kainte receptors did not result in an overt change in the density of neurons expressing these markers in the motor cortex (**Figure S3)** or their distribution across cortical layers I- IV (**Figure S4)**) at any of the timepoints examined. As a subset of neurons in the motor cortex project axons into the corpus callosum, we also analysed transcallosal axon density in the medial corpus callosum of WT and *Grik4^-/-^* mice by TEM (**Figure 6 a**-**d**; **Figure S5**). At P60, axon density was equivalent in WT and *Grik4^-/-^* mice **(Figure S5**). However, unlike WT mice, the *Grik4^-/-^*mice experienced an age-related decline in axon density (**Figure S5**), such that by P180 axon density was ∼44% less than that of P180 WT mice (**Figure 6c-d)**. This was predominantly due to a reduction in the density of unmyelinated axons (**Figure 6c-d**). Furthermore, a higher proportion of myelinated callosal axons showed ultrastructural signs of degeneration in *Grik4^-/-^* mice (**Figure 6e-j**). When degenerating axons were identified, it was not always possible to ascertain whether the axons were originally myelinated or unmyelinated axons, but these were rare, and the density was unchanged by genotype (**Figure 6i)**. However, *Grik4^-/-^* mice had significantly more myelinated axons with disturbed myelin loops that intruded on the axon and were termed inclusions (Lee *et al*., 2012) (**Figure 6i)**. When the density of axons displaying inclusions was expressed as a percentage of the total myelinated axon density (**Figure 6j)**, they represented ∼6% of myelinated axons in WT mice but ∼20% of myelinated axons in *Grik4^-/-^* mice. Despite the significant axonal loss that occurred over time, and the ongoing signs of axon degeneration, immunohistochemical markers of neurodegeneration, including the expression of amyloid precursor protein (APP) and phosphorylated neurofilament (SMI32) were not elevated in the corpus callosum of P180 *Grik4^-/-^* mice (**Figure 6k-v**). The expression of APP and SMI32 were elevated in the corpus callosum of cuprizone-demyelinated mice, which acted as a positive control for gross axonal damage.

**Figure 6.**
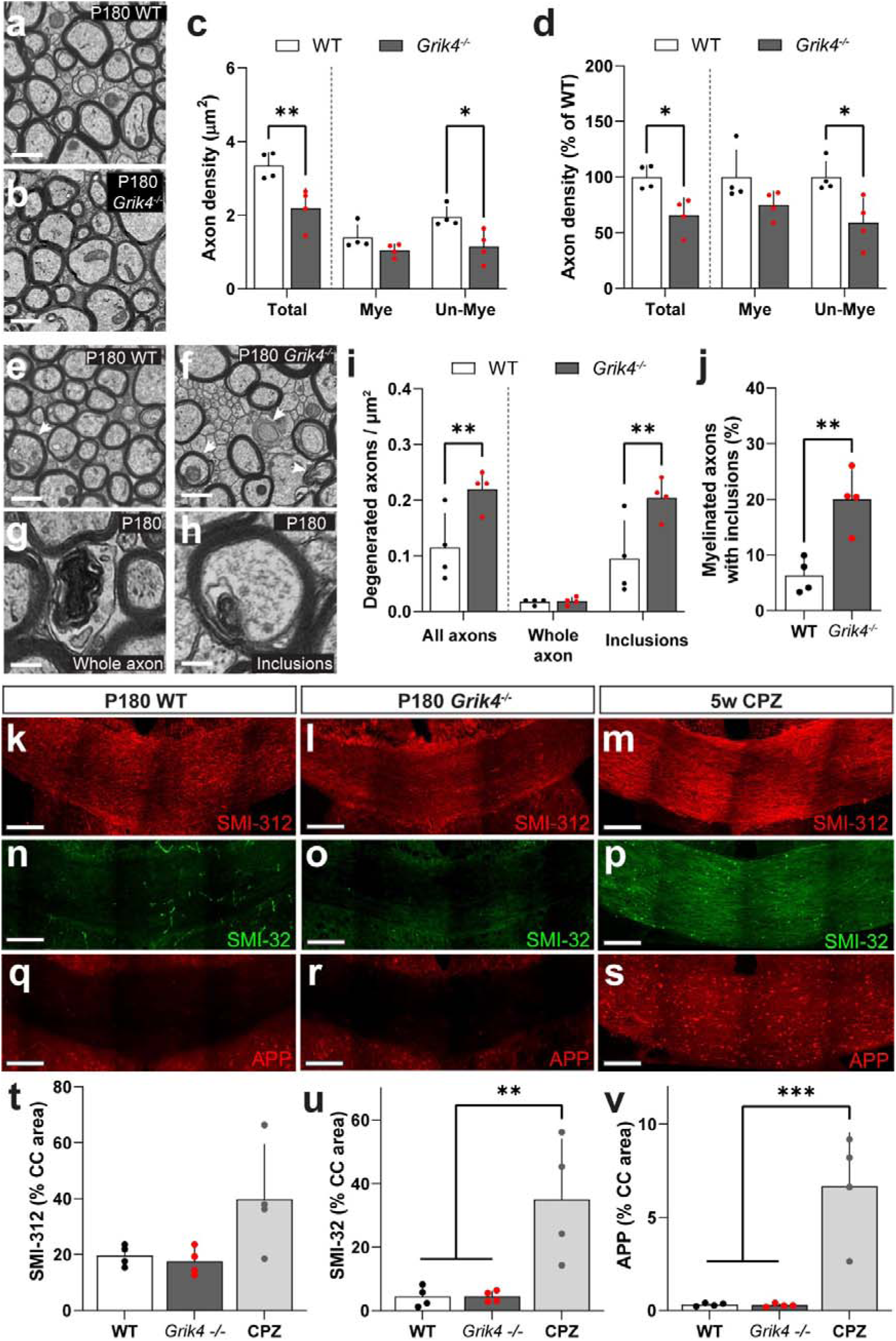
*Grik4^-/-^*mice experience callosal axon loss by P180. **a, b)** Transmission electron micrograph showing myelinated and unmyelinated axons in the medial corpus callosum of a P180 WT (**a**) and *Grik4^-/-^* (**b**) mouse. **c)** The density of total axons as well as myelinated (Mye) and unmyelinated (Un-Mye) axons in the corpus callosum of P180 WT and *Grik4^-/-^* mice [2-way ANOVA: Axon type F(2,18) = 39.91, p<0.0001; Genotype F(1,18) = 25.22, p<0.0001; Interaction F(2,18) = 2.335, p=0.125]. **d)** P180 WT and *Grik4^-/-^* callosal axon density, expressed as a percentage of the average WT density [2-way ANOVA: Axon type F(2,18)=0.4341, p=0.6544; Genotype F(1,18)=21.76, p=0.0002; Interaction F(2,18)=0.4341, p=0.6544]. **e-h)** Transmission electron microscopy images of axons in the corpus callosum of P180 WT and *Grik4^-/-^*mice highlighting an example of whole axon degeneration (**g**) and an inclusion within a myelinated axon (**h**). White arrow heads indicate myelinated axons with inclusions. **i)** Density of callosal axons showing signs of degeneration, and density of axons showing whole axon degeneration or inclusions [2-way ANOVA: Degeneration type F(2,19) = 28.93, p<0.0001; Genotype F(1,18)=16.78, p=0.0007; Interaction F(2,18)=4.071, p=0.035]. **j)** The proportion of myelinated callosal axons with inclusions (% of myelinated axons) [Unpaired t-test: t = 4.428, df = 6, p=0.004]. **k-m)** Images of the corpus callosum of P180 WT, *Grik4^-/-^* mice and WT mice that received 5 weeks of 0.2% cuprizone (CPZ) stained to detect neurofilament SMI-312 (red). **n-p)** Images of the corpus callosum of P180 WT, *Grik4^-/-^* mice and CPZ mice stained to detect phosphorylated neurofilament SMI-32 (green). **q-s)** Images of the corpus callosum of P180 WT, *Grik4^-/-^* mice and CPZ mice stained to detect amyloid precursor protein (APP; red). **t)** Quantification of the area of the corpus callosum (CC) covered by SMI312 labelling (%) [1-way ANOVA F(2, 9)=9.818 p=0.0055]. **u**) Quantification of the area of the corpus callosum (CC) covered by SMI-32 labelling (%) [1-way ANOVA F(2,9) = 4.217, p = 0.051]. **v**) Quantification of the area of the corpus callosum (CC) covered by APP labelling (%) [1-way ANOVA: F(2, 9)=19.39, p=0.0005]. Data presented as mean ± SD for n=4 mice per group. Bonferroni post-test outcomes: * p<0.05, ** p<0.01 or *** p<0.001. Scale bars represent 1µm (**a, b, e, f**); 0.3µm (**g, h**) or 0.1mm (**k-s**).

### The amplitude of callosal CAPs is reduced in Grik4^-/-^ mice

To evaluate the extent to which action potential conduction is impaired in *Grik4^-/-^* mice, we assessed the speed and signal amplitude of APs transmitted along the callosal fibres by measuring the corpus callosum CAP in brain slices from P180 mice. The latency of the peaks corresponding to myelinated and unmyelinated axon fibres was measured over 4 distances of the medial corpus callosum (see schematic in **Figure 7a**). The peak latency was used to determine the conduction velocity of the CAP carried by those nerve fibres. There was no change in the conduction velocity of either myelinated or unmyelinated axon fibres (**Figure 7b-d**). Nor was there a genotype induced change in the half-width of the myelinated or unmyelinated axon peaks (measured at an electrode distance of 1 mm), indicating that the spread of the speed of the fibres contributing to the CAP peaks did not differ in GluK4 functional knockouts (**Figure 7e**, see also **Figure S6** in a second cohort of mice). The refractoriness of the signal amplitude was also unaffected by genotype, indicating that ability of axons to recover after AP conduction was unaffected by the loss of functional GluK4-containing kainate receptors (**Figure S6**).

**Figure 7.**
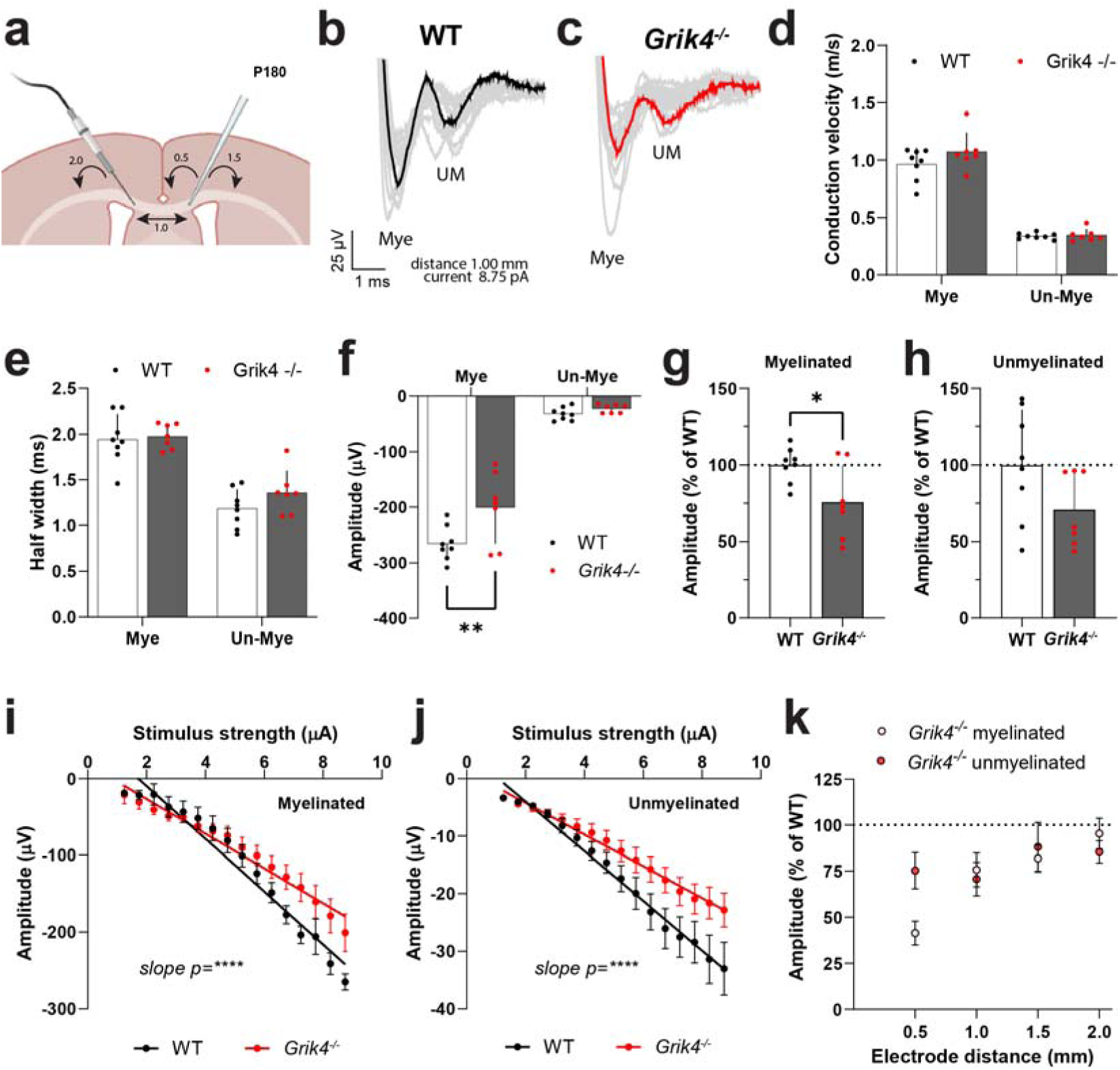
The callosal compound action potential (CAP) is reduced in amplitude but has a normal conduction velocity in *Grik4^-/-^* mice. **a)** Schematic of stimulating and recording electrode placement in the medial corpus callosum in acute brain slices of P180 of WT and *Grik4^-/-^* mice. **b, c)** Voltage traces (grey) from a WT (black) or *Grik4^-/-^* (red) mouse showing the myelinated (Mye; first peak) and unmyelinated (UM; second peak) peaks of the CAP at a maximum stimulus response (8.75 pA) and at an electrode distance of 1 mm. **d)** Conduction velocity of myelinated and unmyelinated axon fibres (average per mouse) [2-way ANOVA: Peak type F(1,26) = 282.2, p<0.0001; Genotype F(1,26)=2.091, p=0.1601; Interaction F(1,26)=1.520, p=0.2287]. **e)** The half width of the CAP myelinated and unmyelinated peaks [2-way ANOVA: Peak type F(1,26) = 72.03, p<0.0001; Genotype F(1,26)=1.613, p=0.2153; Interaction F(1,26)=0.7630, p=0.3904]. **f)** The average amplitude of the CAP myelinated and unmyelinated peaks [2-way ANOVA: Peak type F(1,26) = 250.9, p<0.0001; Genotype F(1,26) = 8.114, p=0.0085; Interaction F(1,26) = 4.498, p=0.044]. **g)** CAP myelinated peak amplitude as a percentage of average WT amplitude (%) [unpaired t-test: t(13)=2.536, p=0.025]. **h)** CAP unmyelinated peak amplitude as a percentage of average WT amplitude (%) [unpaired t-test: t(13)=1.821, p=0.092]. **i)** A plot of stimulus strength versus myelinated peak amplitude (electrode distance 1 mm) [Linear regression slope F(1,236) = 28.99, p<0.0001]. **j)** A plot of stimulus strength versus unmyelinated peak amplitude (electrode distance 1 mm) [Linear regression slope F(1,236) = 21.03, p<0.0001]. **k)** A plot of amplitude (as a percent of WT) versus electrode distance. Data presented as mean ± SD (d-h) or mean ± SEM (i-k). Bonferroni, t-test and linear regression outcomes: * p<0.05, ** p<0.01, **** p < 0.0001 for data from n=7-8 mice per group.

However, when the amplitude of the myelinated CAP peak (electrode distance 1 mm) was measured, it was ∼25% smaller in *Grik4^-/-^*than WT mice (**Figure 7f, g, i**). While the average amplitude of the unmyelinated fibre peak was also ∼30% smaller in *Grik4^-/-^* mice relative to WT mice at this distance, it did not reach statistical significance (**Figure 7f, h, j**). When repeated in a second cohort of mice, both the myelinated and the unmyelinated peaks were significantly smaller in *Grik4^-/-^* compared to WT mice (**Figure S6**). The relative difference in amplitude between *Grik4^-/-^* and WT mice did not increase with increasing distance between the electrodes (**Figure 7k**). These data are consistent with fewer axons contributing to the CAP (i.e. fewer callosal axons), rather than an increased risk of AP failure over longer distances.

### Genes involved in synapse regulation and neuroprotection are differentially expressed in Grik4^-/-^ mice

To gain a broader insight in the mechanisms that lead to behavioural deficits, axon degeneration, and impaired synaptic plasticity in aged *Grik4^-/-^*mice we isolated the hippocampus, as it is a large brain region with the highest level of expression of Gluk4-containing kainate receptors, from aged (>P300) WT and *Grik4^-/-^* mice. At this age *Grik4^-/-^* mice still had an impaired performance on the beam walk test when compared to WT mice (**Figure 8a**). We performed bulk RNA-sequencing and identified 22 differentially expressed genes (DEGs) (**Figure 8b-c**). The list of DEGs included: *Cryab* (LFC = - 1.12; p_adjusted_ = 8.6x10^-25^), which encodes a neuroprotective heat shock protein, αB-crystallin, which is highly expressed by oligodendrocytes and its overexpression can improve motor function in mouse disease models (Hong *et al*., 2017; Oliveira *et al*., 2016); *Gstp2* (LFC = 2.93; p_adjusted_ = 1.4x10^-8^), a gene implicated in cortical neuritogenesis (Liu *et al*., 2021); *Foxg1* (LFC = 2.31; p_adjusted_ = 6.4x10^-9^), a known regulator of cortical circuit development that influences excitatory and inhibitory balance (Hou *et al*., 2020); and three ion channel genes: *Kcnh3* (LFC = 1.89; p_adjusted_ = 4.6x10^-6^; encodes Kv12.2), a known downstream target of *Foxg1* (Vezzali *et al*., 2016) and putative therapeutic target for hyperactivity disorder (Takahashi *et al*., 2018), *Kcnj4* (LFC = 1.45; p_adjusted_ = 4.5x10^-3^), which encodes Kir2.3, an inwardly rectifying potassium channel which localises at the postsynaptic membrane of excitatory synapses and can regulate dendritic morphology (Cazorla *et al*., 2012; Inanobe *et al*., 2002; Trimmer, 2015), and *Scn2b* (-0.40 = 1.89; p_adjusted_ = 1.7x10^-2^), a voltage gated sodium channel beta subunit that modulates CAP amplitude (Chen *et al*., 2002). As the RNA sequencing data suggested that there was a Log_2_ fold change of greater than 1 or -1 in the expression of *Kcnh3*, *Kcnj4* and *Cryab*, and commercial antibodies existed to detect the corresponding proteins (Kv12.2, Kir2.3 and αB-crystallin), we performed Western blots to determine whether gene expression changes were accompanied by protein expression changes in hippocampal lysate from younger (P60) WT and *Grik4^-/-^* mice. We determined that αB-crystallin was significantly downregulated in young adulthood (**Figure 8d-e**), while no change in Kv12.2 or Kir2.3 expression could be detected at this age (**Figure 8f-i**).

**Figure 8:**
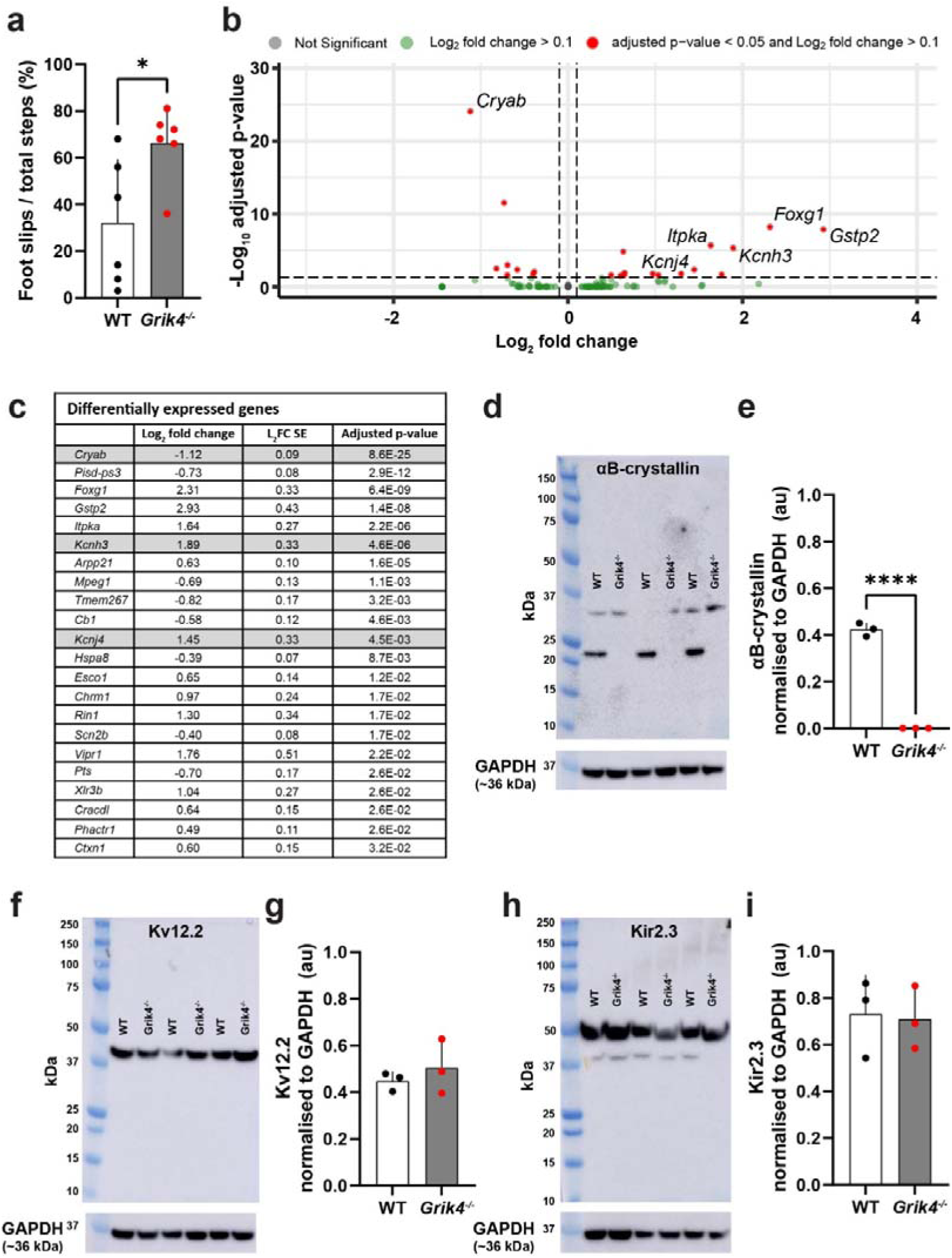
Gene expression differs in the hippocampus of WT and *Grik4^-/-^* mice. **a)** Quantification of foot slips in the beam walk task for WT and *Grik4^-/-^* mice (>P300) [Unpaired t-test: t(10) = 2.66, p=0.024]. **b)** Volcano plot of differentially expressed genes in the hippocampi of WT and *Grik4^-/-^* mice. Differentially expressed genes with an adjusted p-value <0.05 are shown in red. **c)** Table summarising the differentially expressed genes, the Log_2_ fold change and standard error (L_2_FC SE), and adjusted p-values. Grey highlight denotes genes selected for further analysis. **d)** Image of Western blot gel detecting αB-crystallin in hippocampal protein lysates from P60 WT and *Grik4^-/-^* mice (expected size ∼20 kDa). **e**) Quantification of αB-crystallin expression, relative to GAPDH expression [Unpaired t-test: t(4) = 25.68, p <0.0001]. **f, g)** Western blot detection and quantification of Kv12.2 (*KcnH3*) in hippocampal protein lysates from P60 WT and *Grik4^-/-^* mice. We did not observe the expected 100 kDa band - only an ∼40 kDa band, which was unchanged by genotype [unpaired t-test t(4)=0.7733, p = 0.483]. **h-i)** Western blot detection and quantification of Kir2.3 (*Kcnj4*; expected size 49 kDa) was unchanged by genotype [unpaired t-test t(4)=0.1852, p=0.8621]. Footslip and Western blot data are expressed as mean ± SD and t-test results denoted by: *p<0.05 or ****=p<0.0001 for n = 6 mice per genotype (Beamwalk analysis) or n=3 mice per genotype (all other analyses).

## Discussion

### Do GluK4-containing kainate receptors modulate synapse development in the mouse motor cortex?

By P60, layer V pyramidal neurons in the motor cortex of *Grik4^-/-^*mice receive fewer excitatory synaptic inputs than those of WT mice, and basal dendritic spine density is reduced (**Figure 4** and **5**). This could result from the loss of kainate receptor localisation or the loss of ion entry through the affected channels. High affinity kainate receptor subunits (GluK4 or GluK5) are important for targeting kainate receptor subunits to axonal (Vesikansa *et al*., 2012) and synaptic (Palacios-Filardo *et al*., 2016) locations. In *Grik4^-/-^Grik5^-/-^*mice, receptors containing GluK1-3 are no longer targeted to postsynaptic densities or presynaptic active zones in the hippocampus, and kainate-mediated EPSCs cannot be measured at mossy fibre synapses (Fernandes *et al*., 2009); however it is ion entry through kainate receptors that modulate synapse density (Sakha *et al*., 2016).

As we detected a significant loss of thin spines (**Figure 4** and **5**), which are highly plastic and dynamic (Pchitskaya and Bezprozvanny, 2020), Gluk4-containing kainate receptors could regulate excitatory synapse generation and/or turnover in the motor cortex. However, the effect may vary with neuron type. For example, the germline deletion of all five kainate receptor subunits reduces mEPSC frequency and spine density in dorsal striatum medium spinal neurons, but not in CA3 pyramidal neurons (Xu *et al*., 2017). Additionally, the systemic pharmacological antagonism of kainate receptors (UBP-302) during the neonatal period (P1-P15) decreases the density of dendritic spines on CA1 pyramidal neurons, however, this developmental manipulation was associated with an increase in thin and decrease in mushroom spines (Pinzon-Parra *et al*., 2022) – a phenotype different from our observation in the motor cortex. It is interesting to note that the selective deletion of *Grik1* from the basolateral amygdala of P2 mice also reduces mEPSC frequency and spine density in neurons of the centrolateral amygdala by P21 (Ryazantseva *et al*., 2020), however, As *Grik1* is downregulated in the early postnatal period, so that its conditional deletion at P14 no longer alters EPSC frequency or spine density (Ryazantseva *et al*., 2020). From the limited data available, *Grik4* expression in the mouse may instead increase from early development to adulthood in the motor cortex (Hadzic *et al*., 2017; Yue *et al*., 2014), suggesting that GluK4 regulates synapse density in the mouse motor cortex throughout development and adulthood.

The impact of kainate receptor signalling on synapse density appears to differ with neuron type and brain region, however, the receptors would be expected to modulate some common downstream effectors. Our RNA sequencing comparison of gene expression in the hippocampus of WT and *Grik4^-/-^* mice determined that genes capable of regulating synapse number were upregulated in *Grik4^-/-^*mice. These included *Itpka*, *Rin1*, *Gstp2*, *Foxg1* and *Kncj4*. *Itpka* encodes inositol-trisphosphate 3-kinase A, which encodes a CNS neuronal protein (Mailleux *et al*., 1993) that regulates F-actin bundling (Johnson and Schell, 2009), dendritic morphology (Johnson and Schell, 2009; Koster *et al*., 2016; Windhorst *et al*., 2012), and spine number and length in mice (Koster *et al*., 2016). *Rin1* encodes Ras and Rab interactor 1, a Ras-GTPase effector protein that is localised to CNS dendrites (Dhaka *et al*., 2003), regulates the endocytosis of AMPA and other receptors, and regulates the stability of synaptic connections, impacting synaptic plasticity and memory formation (Bliss *et al*., 2010; Deininger *et al*., 2008; Sziber *et al*., 2017). *Gstp2*, encodes glutathione S-transferase Pi 2, which regulates dendritic stability and neurite number (Liu *et al*., 2021). *Foxg1*, encodes a transcription factor that is critical for cortical development (Hou *et al*., 2020), however, its conditional deletion from neurons (*Camk2a* promoter) in P60 mice impairs memory and fear conditioning, and reduces Schaffer-collateral long-term potentiation, coincident with a reduction in CA1 dendritic arborisation and spine density (Yu *et al*., 2019). Lastly, *Kcnj4* encodes the inwardly rectifying potassium channel, Kir2.3, which is implicated in regulating dendritic morphology, as a reduction in Kir2.3 protein is associated with the D2 receptor mediated decrease in the dendritic complexity of dorsal striatal medium spiny neurons (Cazorla *et al*., 2012). The capacity for kainate receptor signalling to modulate synapses may explain previous associations between kainate receptor gene variants, including variants in *Grik4* (Griswold *et al*., 2012; Hu *et al*., 2022; Nisar *et al*., 2022) and the development of neurodevelopmental disorders associated with altered synaptic signalling including schizophrenia (Howes and Onwordi, 2023) and autism (Carroll *et al*., 2021).

### Are Gluk4-containing kainate receptors neuroprotective in the healthy adult mouse brain?

*Grik4^-/-^* mice are hyperactive, show less evidence of depressive- and anxiety-like behaviours in young adulthood (**Figure 2**; Arora *et al*., 2018; Catches *et al*., 2012; Lowry *et al*., 2013) and while P45 *Grik4^-/-^* mice complete the beam crossing test as well as WT mice, their balance and coordination rapidly deteriorate over the following months. By P120 *Grik4^-/-^* mice experience ∼5 times as many hindlimb foot-slips as WT mice (**Figure 2**). This progressive loss of motor coordination temporally correlates with excitatory axon loss. At P60, when mice were hyperactive and had an altered gait, axon density was normal in the corpus callosum of *Grik4^-/-^* mice (**Figures 2 and S5**). However, by P180, when motor coordination was significantly impaired, *Grik4^-/-^* mice had experienced a ∼44% decrease in total callosal axon density, and a higher density of myelinated axons showed ultrastructural signs of neurodegeneration (**Figure 6)**. These changes to callosal axons were associated with a significant reduction in the amplitude of the myelinated and unmyelinated peaks of the callosal compound action potential (**Figure 7 and S6)** and suggest that signalling via Gluk4-containing kainate receptors protects against axon degeneration in the adult mouse brain.

Kainate receptors protecting against neurodegeneration maybe unexpected, given their established role as mediators of excitotoxic injury (Chalupnik and Szymanska, 2023). Indeed, mice that lack functional Gluk4-kainate receptors experience less hippocampal neurodegeneration following an ischemic stroke (Lowry *et al*., 2013). However, it is possible that under physiological conditions, kainate receptors maintain neuronal health. The axon loss detected may be a direct consequence of lost GluK4-containing kainate receptor signalling or a secondary consequence of synapse loss, which can precede neurodegeneration in neurological disorders (Wishart *et al*., 2006). In our RNA-sequencing analysis we found that *Grik4^-/-^* was associated with a significant downregulation in *Cryab,* which encodes the heat shock protein αB-Crystallin, also known as HSPB5. Surprisingly, the protein level of αB-Crystallin protein levels was already reduced by P60. As αB-Crystallin acts to prevent the aggregation of improperly folded proteins and inhibits the apoptotic protein caspase-3 it is neuroprotective (Ousman *et al*., 2007). αB-Crystallin expression is induced in response to cellular stressors (Kirbach and Golenhofen, 2011), and ion entry through GluK4-containing kainate receptors maybe an important inducer of neuroprotective αB-Crystallin. As *Rin1* modulates RAS-GTPases (Sziber *et al*., 2017), which have been linked to the development of neurodegenerative diseases including Alzheimer’s or Parkinson’s disease (for a review see Arrazola Sastre *et al*., 2020), this may be another signalling pathway that leads to neurodegeneration in *Grik4^-/-^* mice. Interestingly, other genes that were upregulated in the *Grik4^-/-^* mice, such as *Foxg1* (Dastidar *et al*., 2011) and *GstP2* (which inhibits JNK signalling (Adler *et al*., 1999; Borsello and Forloni, 2007)) are neuroprotective – and may represent a protective compensatory mechanism induced in response to neurodegeneration.

## Acknowledgments

We thank Anis Contractor (Northwestern University, United States of America) for providing the *Grik4* functional knockout mice. This research was supported by grants from the National Health and Medical Research Council of Australia (NHMRC: 1139041), the Medical Research Future Fund (EPCD00008), the Royal Hobart Hospital research foundation (P0025426), MS Research Australia (16-105; 17-007; 0000000017), and the Australian Research Council (DP180101494; DP220100100). RR, REP and AF were supported by scholarships from the Menzies Institute for Medical Research at the University of Tasmania. CLC and KM were supported by fellowships from MS Research Australia (15-054 and 19-0696). KAP was supported by a fellowship from the National Health and Medical Research Council (NHMRC: 1139180). KMY was supported by fellowships from MS Australia (17-0223; 21-3023). NBB was supported by a fellowship from MS Australia (22-4-097). The funding sources had no role in the study design; collection, analysis or interpretation of the data; writing of the manuscript, or the publication process.

## Data and materials availability

The use of all transgenic mouse lines complies with the conditions of their materials transfer agreements. Individual data points are displayed on the graphs and all source data underlying the graphs are available in Supplementary Data 1. The associated raw image files are a research resource that will be made available by the corresponding author upon reasonable request. The RNA-Seq data, including raw fastq reads, are deposited in NCBI (https://www.ncbi.nlm.nih.gov/) under BioProject ID: PRJNA1076653 (https://www.ncbi.nlm.nih.gov/bioproject/PRJNA1076653). Reviewers can evaluate the data ahead of public release using this link: https://dataview.ncbi.nlm.nih.gov/object/PRJNA1076653?reviewer=ca7s5j1qjmgolfdv9tokgk829h

## Author contributions

KMY, KAP, JC and RR developed the project. RR, KAP, KMY, KM, JF and AF carried out the experiments. KMY, KAP, CLC, NB, and JC obtained research funding. RR, KAP, NB, JF, KM and AF performed the statistical analyses and generated the figures. KMY, KAP, REP, JC, CLC and WC provided methodological expertise, training, and supervision. KMY, RR and KAP wrote the manuscript. All authors have edited and approved the final manuscript.

