## Supplementary Figures for "Gluk4-containing kainate receptors regulate synaptic communication in the motor cortex and reduce axon degeneration in adult mice"

**Supplementary Material Ricci et al.**

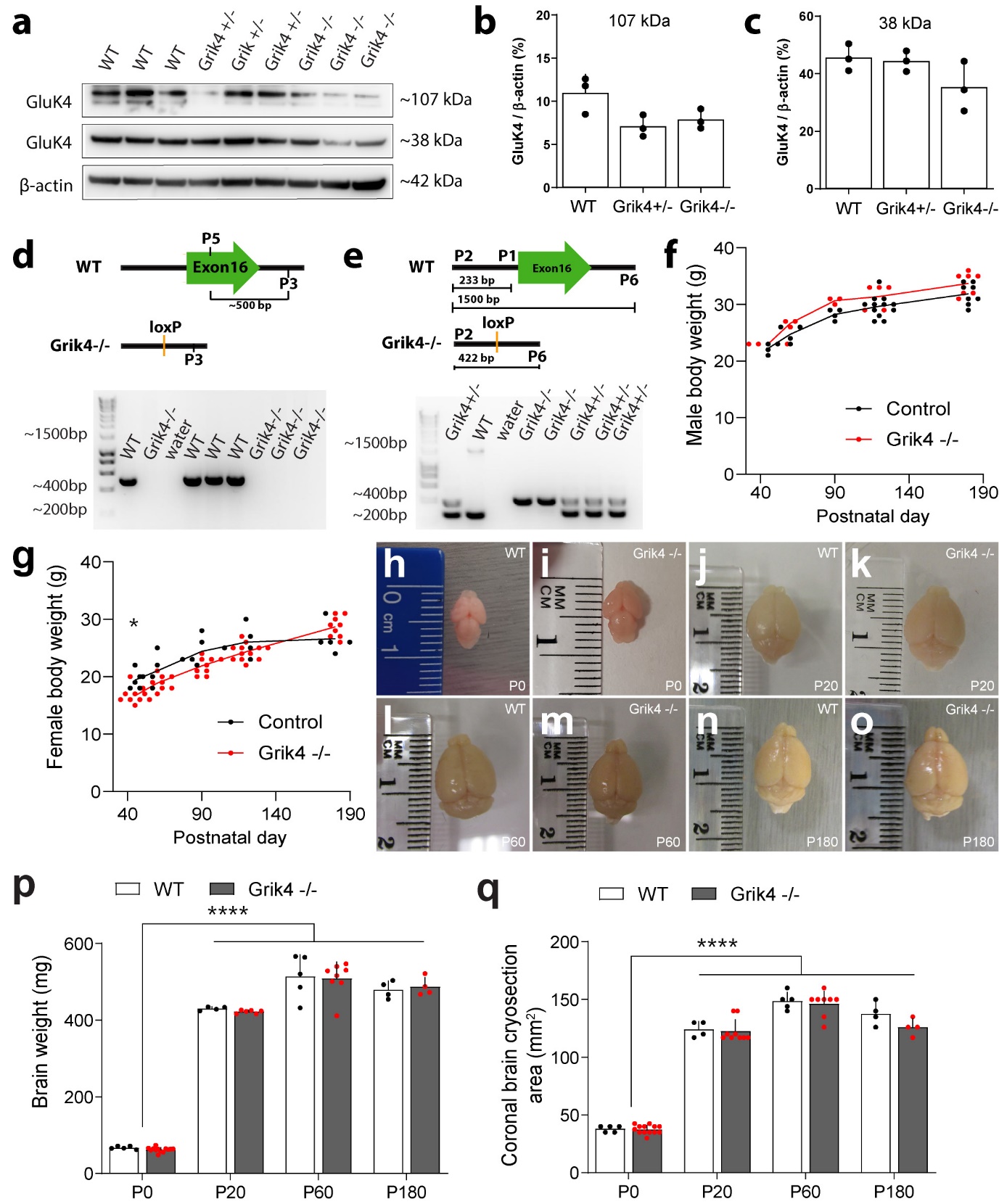

**Figure S1. *Grik4^-/-^* mice lack exon 16 but produce normal levels of GluK4 protein and maintain normal body and brain weight. a)** Western blot detection of GluK4 and β-actin in brain lysates from P5 WT, *Grik4^+/-^* and *Grik4^-/-^* mice. Anti-Gluk4 detected the expected 107 kDa band, but also detected a smaller 38kDa band that has been previously reported (Marrocco et al. 2012) and may correspond to an uncharacterized cleavage product or reflect non-specific binding of the antibody. **c, d)** Quantification of the ~107 kDa Gluk4 protein band, relative to β-actin expression, in brain lysates from P5 WT, *Grik4^+/-^* and *Grik4^-/-^* mice [one way ANOVA: F(2,6) = 4.99, p=0.053]. **d)** Quantification of the ~38 kDa Gluk4 protein band, relative to β-actin expression, in brain lysates from P5 WT, *Grik4^+/-^* and *Grik4^-/-^* mice [one way ANOVA: F(2,6) = 2.59, p=0.155]. **e)** Gel photo of PCR products resulting from the amplification of genomic DNA from P20 WT and *Grik4^-/-^* mice using the P5 primer (AGGGCCGGTGTAATCTCCTG) which binds within exon 16 of *Grik4* and the P3 primer (CTCTGTACGCAGACCCCAGAG) downstream of exon 16. This amplifies a ~400bp fragment in WT DNA and no amplification is seen from *Grik4^-/-^* DNA. **f, g)** Gel photo of PCR products resulting from the amplification of genomic DNA from P20 WT and *Grik4^-/-^* mice using the P1, P2 and P6 primers (sequence in materials and methods). This PCR produces an ~233bp fragment corresponding to WT Grik4 and an ~422bp fragment corresponding to the *Grik4* gene that lacks exon 16. **h)** Body weight of male WT and *Grik4^-/-^* at P0, P20, P60 and P180 [2-way ANOVA: Age F(4,43) = 52.80, p<0.0001; Genotype F(1,43) = 11.50, p=0.0015; Interaction F(4,43)=0.2255, p = 0.923]. Bonferroni multiple comparison indicates no significant effect of genotype. **i)** Body weight of female WT and *Grik4^-/-^* at P0, P20, P60 and P180 [2-way ANOVA: Age F(4,63) = 57.27, p<0.0001; Genotype F(1,63) = 9.321, p=0.003; Interaction F(4,63) = 3.693, p=0.009]. Bonferroni multiple comparison indicates that at P45, female *Grik4^-/-^* mice may be slightly smaller than WT mice, p=0.05]. **j-q)** Example images of brains from P0, P20, P60 and 180 WT and *Grik4^-/-^* mice. **r)** Brain weight at P0, P20, P60 and P180 for WT and *Grik4^-/-^* mice [2-way ANOVA: Age F(3,41) = 778.9, p<0.001; Genotype F(1,41) = 0.095, p = 0.76; Interaction F(3,41) = 0.15, p=0.93]. **j)** Average area (in mm^2^) of brain coronal cryosections from P0, P20, P60 and P180. In adulthood cryosections were collected at ~Bregma 3. [2-way ANOVA: Age F(3,44) = 550. P<0.0001; Genotype F(1,44) = 3.025, p=0.089; Interaction F (3,44) = 1.033, p=0.387]. Data are presented as mean ± SD. * p<0.05 Bonferroni; **** p<0.0001 2-way ANOVA age effect. n=3 mice per group for Western blot analyses; 5-10 mice per group for weight analyses.

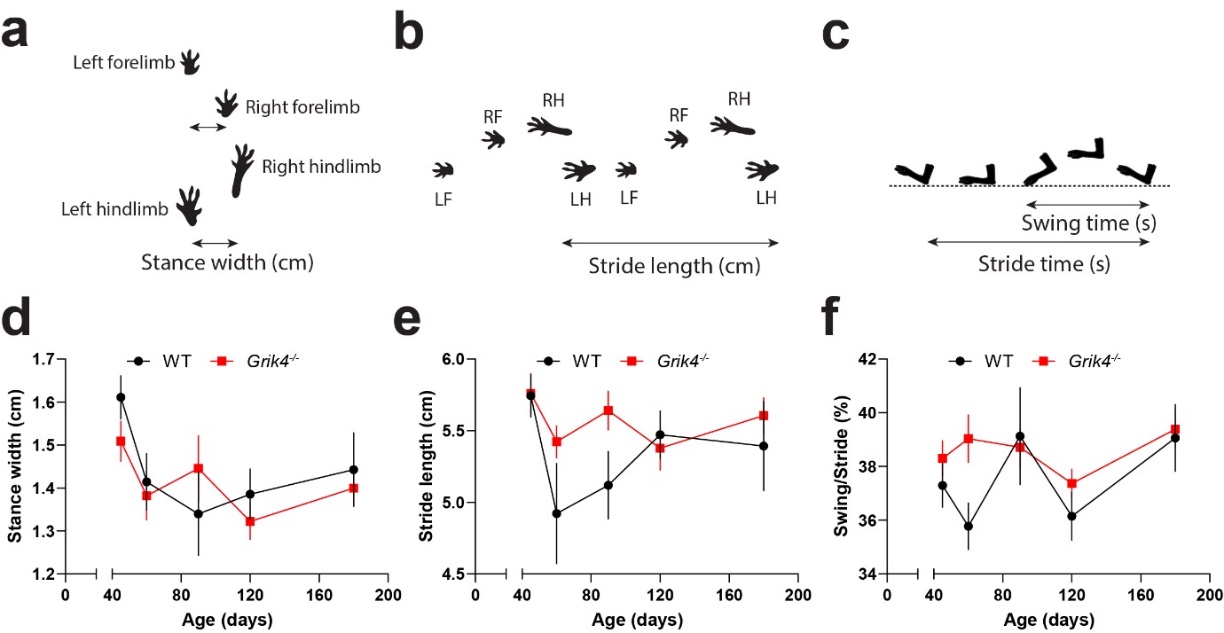

### Figure S2. Forelimb gait parameters are normal in *Grik4^-/-^* mice.

**a-c)** Schematic diagrams illustrating the measurement of stance width, stride length and swing time. WT and *Grik4^-/-^* mice ran on the Digigait at 22cm/s and repeated measurements were taken between P45 and P180 to determine: **d)** forelimb stance width [2-way ANOVA: Age F(4,77)=3.481, p=0.0115; Genotype F(1,77)=0.45, p=0.504, Interaction F(4,77)=0.702, p=0.59]; **e)** forelimb stride length [2-way ANOVA: Age F(4,77)=2.782, p=0.032; Genotype F(1,77) = 3.752, p=0.056; Interaction F(4,77) = 1.7, p = 0.376], or **f)** forelimb swing time, as a percentage of stride time [2-way ANOVA: Age F(4,77) = 2.531, p=0.047; Genotype F(1,77) = 3.56, p=0.063; Interaction F(4,77) = 1.134, p=0.347]. Data presented as mean ± SEM for n=5-11 mice per group.

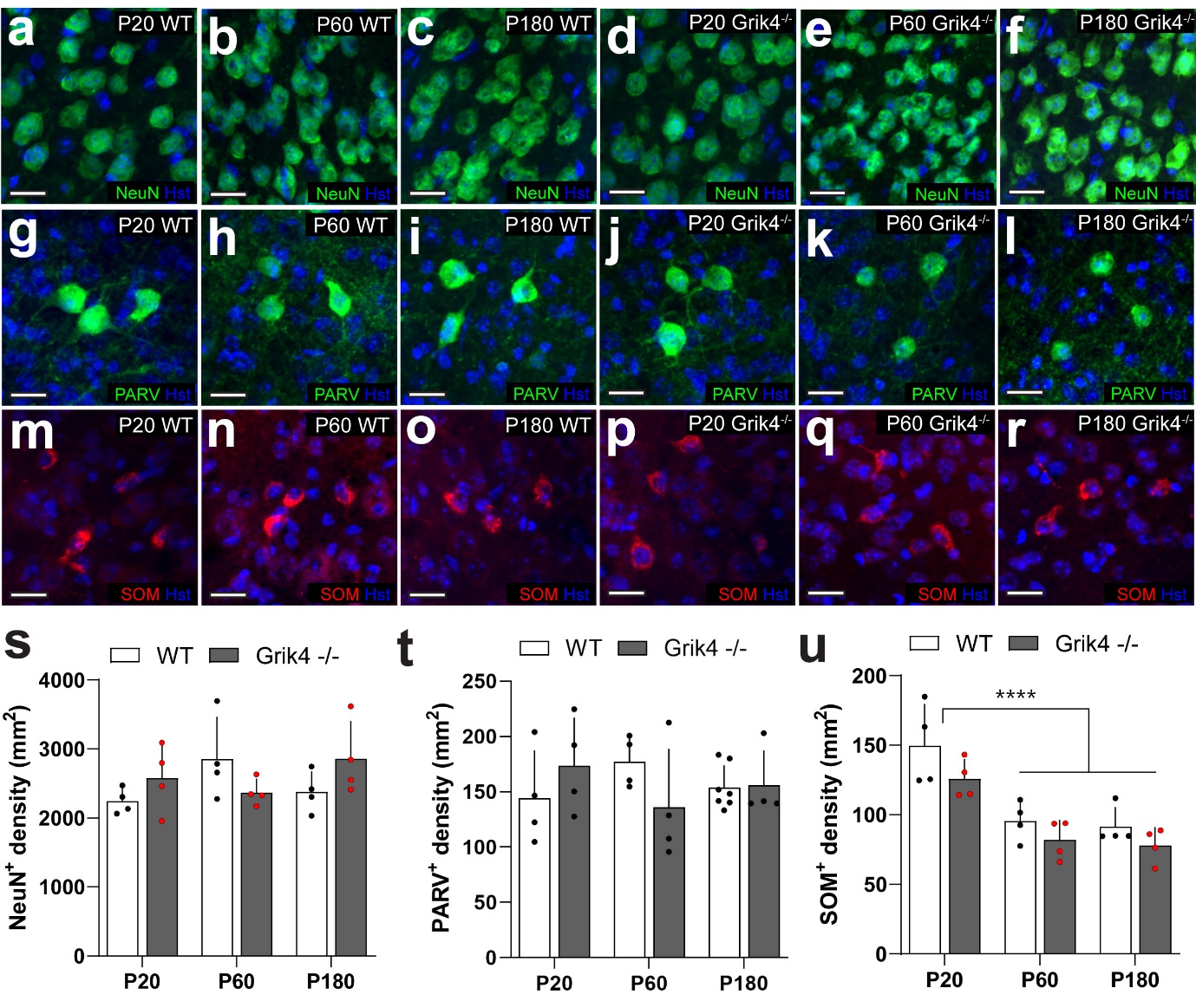

**Figure S3. Neuron and interneuron density in the primary motor cortex is not affected by GluK4 knock-out.** Example images of **a-f**) total neurons (NeuN+ cells), **g-l**) parvalbumin positive (PARV+) interneurons, and **m-r)** somatostatin positive (SOM+) interneurons in the primary motor cortex of WT and *Grik4^-/-^* mice at postnatal day 20, 60 and 180. **s)** The density of NeuN+ cells was unchanged in the *Grik4^-/-^* mice [2-way ANOVA: Age F(2,18)=0.6344, p=0.5417; Genotype F(1,18)=0.4215, p=0.5244; Interaction F(2,18)=3.114, p=0.0690] **t)** The density of PARV+ cells was unchanged in the primary motor cortex of *Grik4-/-* mice [2-way ANOVA: Age F(2,21)=0.02978, p=0.9707; Genotype F(1,21)=0.05689, p=0.8138; Interaction F(2,21)=1.994, p=0.1611] **u)** The density of SOM+ cells decreased in both WT and *Grik4^-/-^* mice after P20, but there was no difference between WT and *Grik4^-/-^* mice [2-way ANOVA; Age F(2,18)=23.02 p<0.0001, Genotype F(1,18)=5.761 p=0.0274, Interaction F(2,18)=0.2326 p=0.7948]. Values represent mean ± SD, ****p = < 0.0001 2-way ANOVA main effect, n=4 animals for all groups.

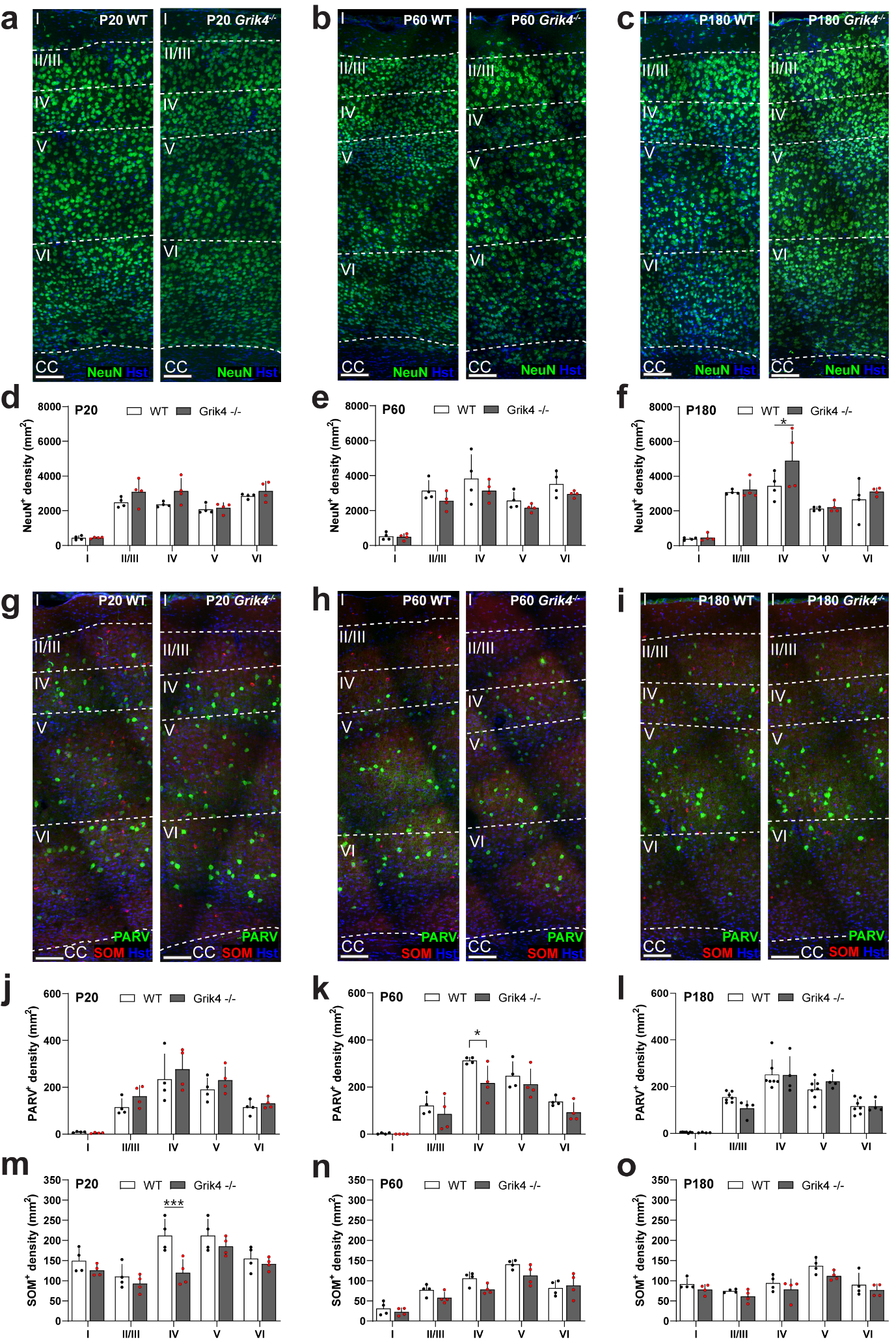

**Figure S4. GluK4 functional deletion does not affect neuron or interneuron cell density in the layers of the primary motor cortex. a-c)** Example images of neurons (NeuN+ cells) in primary motor cortex layers 1 to 6 from WT and *Grik4-/-* mice at postnatal day 20, 60 and 180. **d-f)** Cell density in cortical layers was unchanged in the *Grik4-/-* mice. [2-way ANOVA with Bonferroni’s post-test: P20 Interaction F(4, 30)=1.332 p=0.2810, Layer F(4, 30)=52.81 p<0.0001, Genotype F(1, 30)=7.534 p=0.0101; P60 Interaction F(4, 30)=0.3656 p=0.8311, Layer F(4, 30)=30.63 p<0.0001, Genotype F(1, 30)=5.710 p=0.0234; P180 Interaction F(4, 30)=1.287 p=0.2973, Layer F(4, 30)=29.20 p<0.0001, Genotype F(1, 30)=3.694 p=0.0642]. **g-i)** Example images of parvalbumin (PARV+) or somatostatin (SOM+) positive interneurons in primary motor cortex layers 1 to 6 from WT and *Grik4-/-* mice. **j-o)** *Grik4* functional knock-out does not affect interneuron cells density in the cortical layers in the primary motor cortex of mice at postnatal day 20, 60 and 180. [2-way ANOVA with Bonferroni’s post-test: Parvalbumin P20 Interaction F(4, 30)=0.3128 p=0.8672, Layer F(4, 30)=24.28 p<0.0001, Genotype F(1, 30)=2.701 p=0.1107; P60 Interaction F(4, 30)=1.042 p=0.4020, Layer F(4, 30)=40.34 p<0.0001, Genotype F(1, 30)=8.462 p=0.0068; P180 Interaction F(4, 45)=1.353 p=0.2653, Layer F(4, 45)=54.66 p<0.0001, Genotype F(1, 45)=0.07120 p=0.7908: Somatostatin P20 Interaction F(4, 30)=2.486 p=0.0646, Layer F(4, 30)=12.02 p<0.0001, Genotype F (1, 30)=14.15 p=0.0007; P60 Interaction F(4, 30)=1.347 p=0.2756, Layer F(4, 30)=33.92 p<0.0001, Genotype F(1, 30)=7.378 p=0.0108; P180 Interaction F(4, 30)=0.1582 p=0.9577, Layer F(4, 30)=11.09 p<0.0001, Genotype F(1, 30)=8.192 p=0.0076]. Values represent mean ± SD, *p < 0.05, ***p < 0.001 2-way analysis of variance (ANOVA) with Bonferroni’s post-test, n=4-7 animals for all groups.

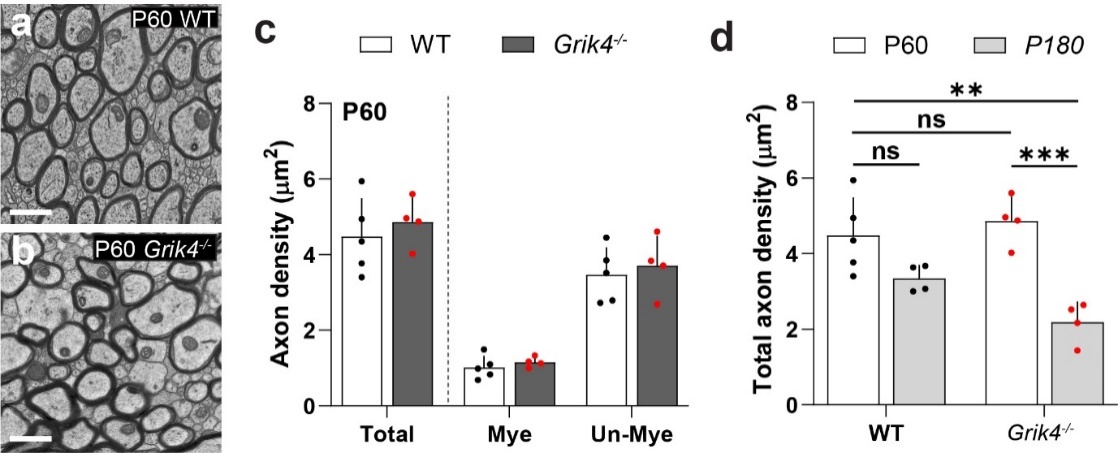

**Figure S5. Axon density is unchanged by genotype at P60 but declines with age in the *Grik4^-/-^* mice. a-b)** Example images of axon ultrastructure in the medial corpus callosum of wild type (WT) and *Grik4^-/-^* mice at postnatal day (P) 60. **c)** Axon density in the corpus callosum at P60 is equivalent in WT and *Grik4^-/-^* mice [2-way ANOVA: Axon type F(2,21)=65.02, p<0.0001; Genotype F(1,21)=0.9362, p=0.3443; Interaction F(2,21)=0.07180, p=0.9309]. **d)** There is an age-related decrease in total axon density between P60 and P180 in *Grik4^-/-^* mice, whereas there is not in WT mice [2-way ANOVA: Genotype F(1,13)=1.243, p=0.2851; Age F(1,13)=30.28, p=0.0001; Interaction F(1,13)=4.928, p=0.0448]. Data is presented as mean ± SD, ** p<0.01, *** p<0.001 Šídák's multiple comparisons test, n = 4-5 mice in each group.

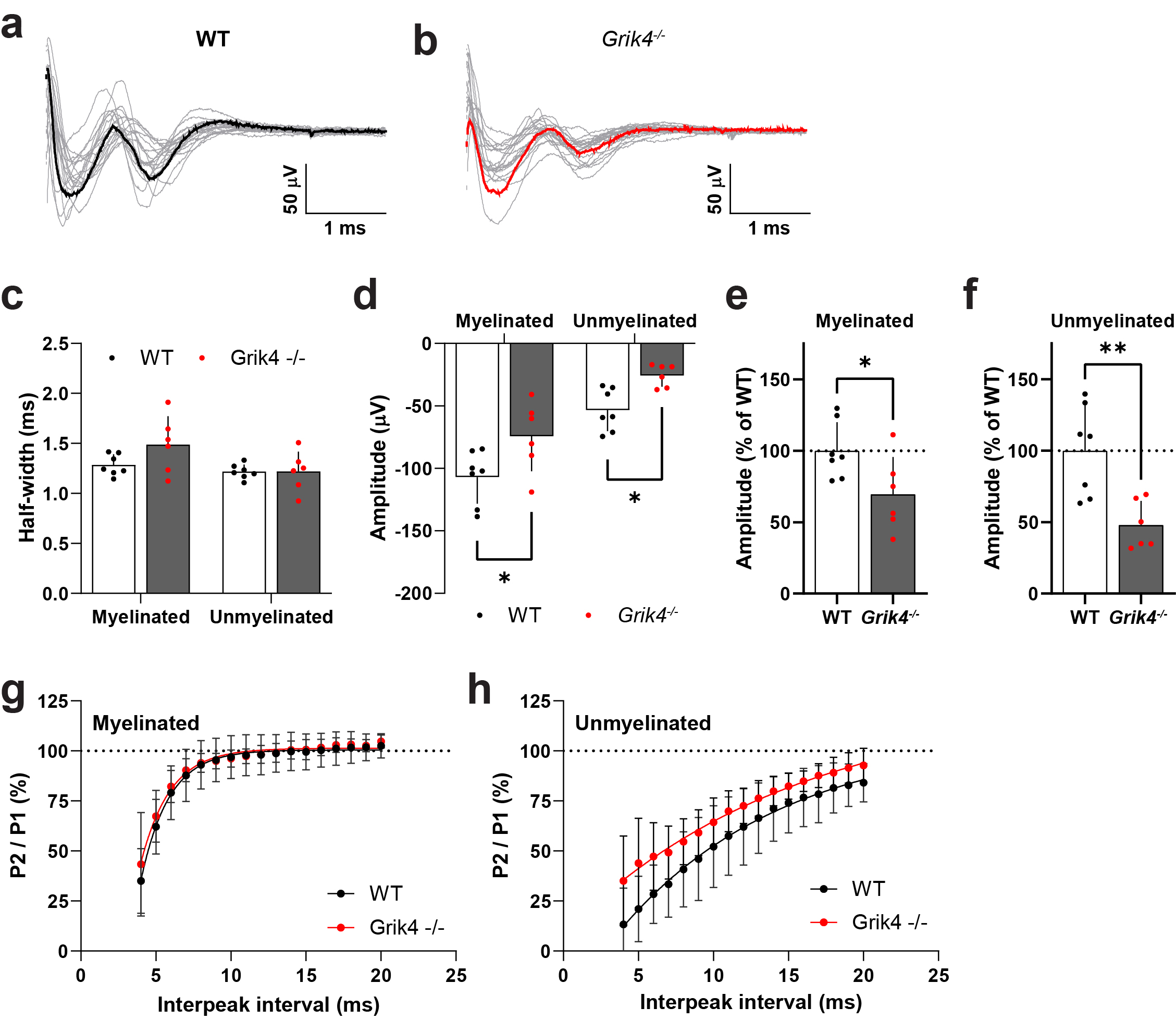

**Figure S6. Gluk4 functional knock-out does not affect refractoriness of the myelinated or unmyelinated CAP peaks. a-b)** Voltage traces (grey) showing the myelinated (first peak) and unmyelinated (second peak) peaks of the CAP in a second cohort of P180 mice (stimulus response 8.75 pA and electrode distance 1 mm). Black (WT) or red (*Grik4^-/-^*) traces highlight an example trace. **c)** Half-width of the peaks for both the myelinated and unmyelinated axon fibres did not differ between WT and *Grik4^-/-^* mice [2-way RM ANOVA: Peak F(1,11)=7.652, p=0.0183; Genotype F(1,11)=1.712, p=0.2174; Interaction F(1,11)=2.768, p=0.1244] **d)** The mean amplitude of the myelinated and unmyelinated peaks was reduced in the *Grik4^-/-^* mice [2-way RM ANOVA: Peak F(1,11)=50.43, p<0.0001; Genotype F(1,11)=12.77, p=0044; Interaction F(1,11)=0.1192, p=0.7364] **e)** When expressed relative to the mean WT myelinated peak amplitude, there was an approximately 30% reduction in myelinated peak amplitude in the *Grik4^-/-^* mice [unpaired t-test t=2.388, p=0.0360] **f)** When expressed relative to the mean WT unmyelinated peak amplitude, there was an approximately 50% reduction in unmyelinated peak amplitude in the *Grik4^-/-^* mice [unpaired t-test t=3.601, p=0.0042]. **g)** The refractoriness of the amplitude of the myelinated peak (second peak divided by the first peak, expressed as a percent) did not differ between WT and *Grik4^-/-^* mice at any of the interpeak time intervals measured [2-way RM ANOVA Time interval F(1.536,16.90)=95.70, p<0.001, Genotype F(1,11)=0.1569, p=0.6996, Interaction F(16,176)=0.4401, p=0.0697]. **h)** The refractoriness of the amplitude of the unmyelinated peak also did not differ by genotype [2-way RM ANOVA: Time interval F(1.676,18.44)=66.54, p<0.0001; Genotype F(1,11)=3.637, p =0.0829; Interaction F(16,176)=0.9033, p=0.05664]. Values represent mean ± SD. Asterixis represent the results of Šídák's multiple comparisons post-hoc test, or unpaired t-tests, as appropriate. *=p<0.05, **=p<0.01. n = 6-7 mice for all groups.
